# Identification of Small Molecule Ligand Binding Sites On and In the ARNT PAS-B Domain

**DOI:** 10.1101/2023.11.03.565595

**Authors:** Xingjian Xu, Joseph Closson, Leandro Pimentel Marcelino, Denize C. Favaro, Marion L. Silvestrini, Riccardo Solazzo, Lillian T. Chong, Kevin H. Gardner

**Author notes:** address correspondence to: Kevin Gardner, Structural Biology Initiative, CUNY Advanced Science Research Center, 85 St. Nicholas Terrace, New York, NY 10031 USA.

## Abstract

Transcription factors are generally challenging to target with small molecule inhibitors due to their structural plasticity and lack of catalytic sites. Notable exceptions include several naturally ligand-regulated transcription factors, including our prior work with the heterodimeric HIF-2 transcription factor which showed that small molecule binding within an internal pocket of the HIF-2α PAS-B domain can disrupt its interactions with its dimerization partner, ARNT. Here, we explore the feasibility of similarly targeting small molecules to the analogous ARNT PAS-B domain itself, potentially opening a promising route to simultaneously modulate several ARNT-mediated signaling pathways. Using solution NMR screening of an in-house fragment library, we previously identified several compounds that bind ARNT PAS-B and, in certain cases, antagonize ARNT association with the TACC3 transcriptional coactivator. However, these ligands have only modest binding affinities, complicating characterization of their binding sites. We address this challenge by combining NMR, MD simulations, and ensemble docking to identify ligand-binding ‘hotspots’ on and within the ARNT PAS-B domain. Our data indicate that the two ARNT/TACC3 inhibitors, KG-548 and KG-655, bind to a β-sheet surface implicated in both HIF-2 dimerization and coactivator recruitment. Furthermore, while KG-548 binds exclusively to the β-sheet surface, KG-655 can additionally bind within a water-accessible internal cavity in ARNT PAS-B. Finally, KG-279, while not a coactivator inhibitor, exemplifies ligands that preferentially bind only to the internal cavity. All three ligands promoted ARNT PAS-B homodimerization, albeit to varying degrees. Taken together, our findings provide a comprehensive overview of ARNT PAS-B ligand-binding sites and may guide the development of more potent coactivator inhibitors for cellular and functional studies.

## Introduction

There has been considerable interest in targeting non-enzymatic protein-protein interfaces for therapeutic purposes (1, 2). Transcription factor complexes are a primary target of this approach, as small molecule disruptors or degraders of such complexes could potentially modulate the aberrant gene expression associated with many diseases. However, with few notable exceptions, transcription factors are perceived as ‘undruggable’ due to their lack of targetable catalytic sites (3). Finding small molecules that can specifically bind to and disrupt the large, flat surfaces involved in protein-protein or protein-DNA interactions remains a significant challenge (4–6). Here, we focus on the characterization of small molecule interactions with a transcription factor subunit, aryl hydrocarbon receptor nuclear translocator (ARNT), as a case study of exploiting structural facets in the development of inhibitors that target transcription factor/coactivator interactions.

ARNT is a basic Helix Loop Helix – Per-ARNT-Sim (bHLH-PAS) transcription factor, so named for its N-terminal DNA binding bHLH motif and two PAS domains (PAS-A and PAS-B) which all mediate binding the requisite DNA and protein components required for function (7). The bHLH-PAS proteins can be broadly divided into the signal-regulated class I and the constitutively expressed class II subunits. ARNT is a class II protein which serves as a ‘universal’ binding partner to several class I bHLH-PAS proteins, including the hypoxia-inducible factors (HIF-α), aryl hydrocarbon receptor (AHR), neuronal PAS proteins (NPAS), and single-minded proteins (SIM) (7). Once formed, these heterodimers control a plethora of biological processes, including hypoxia adaptation, xenobiotic metabolism, and neurogenesis (7).

Two well-characterized ARNT-containing signaling cascades which respond to environmental stimuli are the xenobiotic-sensing AHR and hypoxia-regulated HIF pathways (8) (**Fig. 1**). For the former, AHR regulates the vertebrate xenobiotic response pathway by its binding to a wide variety of small molecules, ranging from pollutants (e.g., 2,3,7,8 – tetrachlorodibenzo-p-dioxin (TCDD, “dioxin”)) to dietary compounds (9–11). In the cytosol, inactivated AHR is bound to and stabilized by the heat shock protein 90 (HSP90) chaperone and other auxiliary proteins (12, 13). Ligand binding to the AHR PAS-B domain triggers a poorly-understood response which includes translocation of cytosolic AHR into the nucleus, where it heterodimerizes with ARNT to form the functional heterodimeric transcription factor that binds to dioxin response elements (DREs) upstream of AHR-controlled genes and initiates their transcription (14). The HIF pathway, on the other hand, mediates cellular adaptation to low oxygen levels (hypoxia) (15–17). Under normoxia, the HIF-α subunits (HIF-1α, 2α, 3α) are rapidly degraded and inactivated through oxygen-dependent post-translational modification pathways; these controls are relieved under hypoxic conditions, allowing the HIF-α subunits to accumulate and dimerize with ARNT in the nucleus (17). The HIF-α/ARNT heterodimer complexes bind to hypoxia response elements (HREs) upstream of HIF-controlled genes, leading to erythropoiesis, angiogenesis, and metabolic remodeling (16–18).

**Fig. 1.**
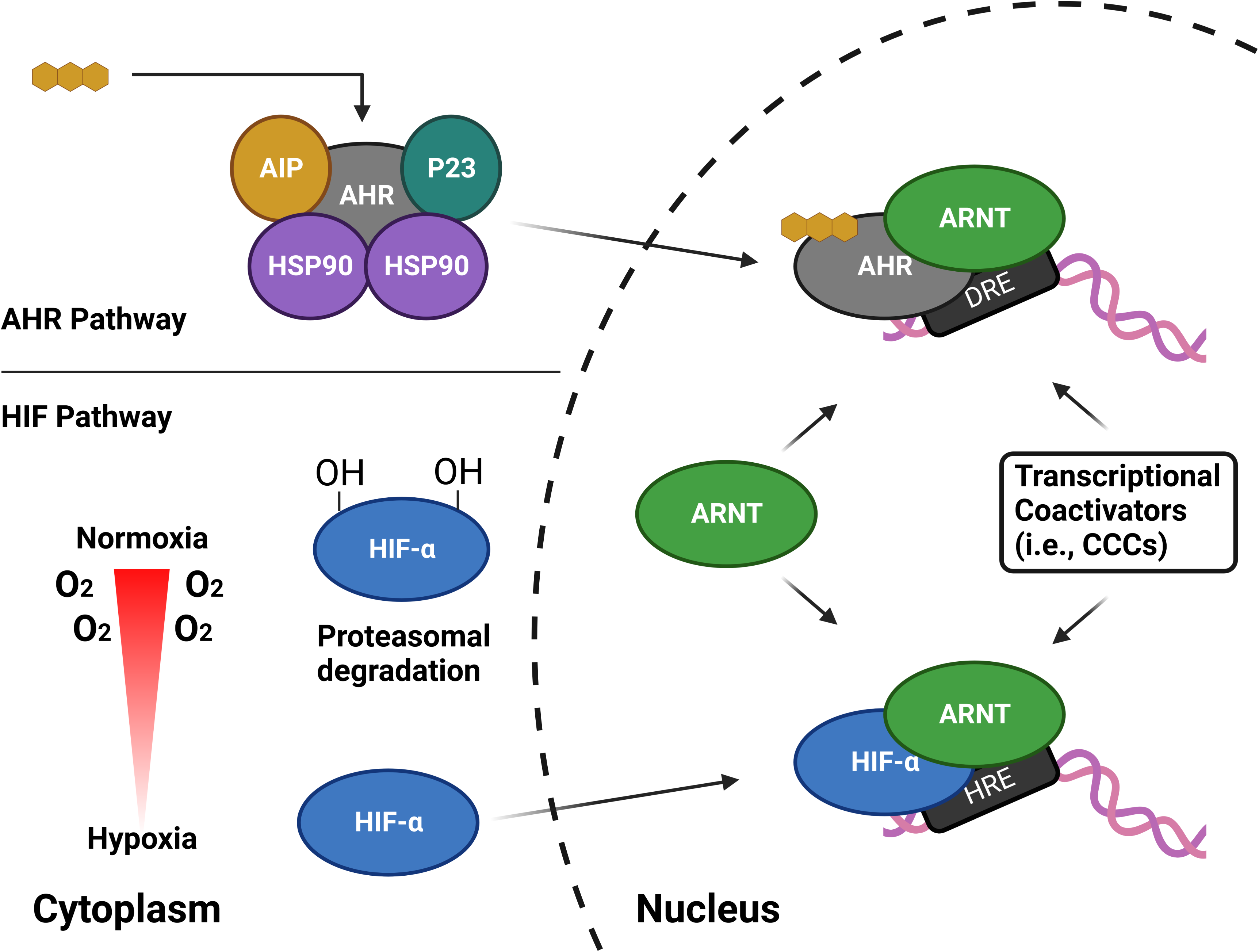
Schematic diagram of the AHR and HIF pathways. In the quiescent state, AHR resides in the cytoplasm in a complex with other proteins, including heat shock protein 90 (Hsp90), aryl hydrocarbon receptor interacting protein (AIP), and p23. Upon ligand binding, AHR translocates to the nucleus and dimerizes with ARNT. HIF activation is oxygen dependent, via O_2_-utilizing hydroxylases. Under normoxic conditions, the HIF-α subunits are rapidly degraded and inactivated by downstream effects of hydroxylation. Under hypoxic conditions, hydroxylation fails to occur, letting HIF-α dimerize with the β-subunit (ARNT) in the nucleus. Both AHR and HIF complexes recruit various coactivators as part of the transcription activation process.

Dysregulation of both the AHR and HIF pathways has been implicated in many diseases. For example, AHR is an important modulator of host-environment interactions and plays critical roles in immune and inflammatory responses (19). Overactivation of AHR can also stimulate tumorigenesis, making it a promising (but still underexplored) therapeutic target for cancer therapies (20, 21). In comparison, anomalous HIF activities may lead to anemia, inflammation, and ischemia (16, 22–25). Persistent HIF activation can also contribute to tumor growth and progression, aiding tumor cell survival and therapy resistance (22, 25, 26). While several strategies targeting different parts of the HIF pathway have been tested, the most efficacious ones focus solely on the HIF-2α isoform (18, 27–31), including the clinically-utilized HIF-2α inhibitor WELIREG (belzutifan) (32–38).

As with many other transcription factors, AHR and HIF rely on a variety of functionally important coactivators to execute their programs (39–42), opening potential other opportunities for small molecule control of their signaling properties. These include a group of proteins known as the coiled-coil coactivators (CCCs) (41, 43–47) which are recruited via the ARNT PAS-B domain and thus establish the ARNT PAS-B/coactivator interaction as a potential target for therapeutic intervention (41, 44, 46). One such CCC, the coiled-coil protein coiled-coil coactivator (CoCoA), has been proposed as a primary coactivator of the AHR/ARNT complex, and interacts with the bHLH-PAS regions of both proteins (45), suggesting that ARNT/CCC interactions could be a point of modulation for both HIF and AHR pathways. Using NMR-based fragment screening and *in vitro* interaction assays, we identified 10 compounds that bound ARNT PAS-B with micromolar affinities, including two compounds (KG-548, KG-655) capable of disrupting its interactions with a fragment of the Transforming Acidic Coiled-Coil Containing Protein 3 (TACC3) CCC protein (48). While characterizing the binding sites of these ligands has been technically challenging, we recently reported an ARNT/KG-548 co-crystal structure showing that KG-548 binds on the surface of ARNT PAS-B (49). Although this binding site is reasonable and appears to be privileged for protein and small molecule interactions (50), validation of this binding mode is essential as binding occurs at a dimeric interface between two ARNT PAS-B molecules that we have not seen stably in solution (49). Further, we have been unable to use a similar crystallographic approach to obtain structures of ARNT PAS-B bound to KG-655 or other compounds, leaving their binding modes unclear.

To address these knowledge gaps, we investigated the binding modes of several small molecules with the ARNT PAS-B domain using solution NMR spectroscopy, weighted ensemble molecular dynamics (WEMD) simulations, and ensemble docking. We used three ligands, KG-548, KG-655, and KG-279, as proof-of-concept small molecules to showcase two ligand binding modes: one on the external side of the PAS β-sheet (consistent with our KG-548/ARNT crystal structure (49)) and an internal cavity previously seen by X-ray diffraction (48). Notably these three compounds differentially preferred binding these sites, likely in correlation to their sizes and chemical features. We also used ensemble docking to determine potential binding poses of KG-655 and KG-279 in the internal cavity in ways that satisfied NOE-based protein/ligand distances, showing marked distortion of the protein domain upon ligand binding. Taken together, this study validates the previously proposed surface binding site for KG-548 and provided the first direct evidence of ligand binding inside the ARNT PAS-B domain cavity, all of which will aid designing more potent ARNT PAS-B/coactivator inhibitors for functional studies.

## Materials and Methods

### Protein expression

Wildtype human ARNT PAS-B (residues 356-470) and point mutants (I364V, I396V, Y456T, I457V, I458V, I364V/I458V) were cloned into the pHis-parallel bacterial expression vector (51). His_6_-tagged ARNT PAS-B proteins were overexpressed in BL21(DE3) *E. coli* (Agilent). Uniformly labeled (U-^13^C,^15^N) proteins were obtained by growing cells in M9 minimal media supplemented with 3 g/L U-^13^C_6_-D-glucose and 1 g/L ^15^NH_4_Cl. To obtain stereospecific NMR assignments of prochiral methyl groups of valines and leucines, a fractional ^13^C-labeling strategy was used with 10% (0.3 g/L) of U-^13^C_6_-D-glucose and 90% (2.7 g/L) of unlabeled D-glucose added to the M9 minimal media (52). For all expressions, cells were induced with 0.5 mM isopropyl-β-D-thiogalactopyranoside (IPTG) at 37 °C after reaching OD_600_ of 0.6 – 0.8. Cells were incubated overnight (16-18 hrs) at 18 °C and harvested by spinning at 4 °C, 4658 g for 30 minutes. Pellets were flash-frozen with liquid nitrogen and stored at –80 °C until purification.

### Protein purification

Protein pellets were resuspended in lysis buffer (50 mM Tris-HCl, pH 7.4 at room temperature (RT), 150 mM NaCl, 10 mM imidazole, 1 mM DTT, 1 mM PMSF), lysed by sonication, and centrifuged at 47850 g for 45 minutes. Purification started with an initial affinity chromatography step using a gravity column with nickel-Sepharose High-Performance resin or HisTrap HP (Cytiva). First, protein-bound columns were washed with the wash buffer (50 mM Tris-HCl, pH 7.4 at RT, 150 mM NaCl, 1 mM DTT) with increasing concentration of imidazole (up to 150 mM). Next, His_6_-tagged proteins were eluted with the elution buffer (wash buffer + 500 mM imidazole) and subsequently diluted 1:10 with cleavage buffer (50 mM Tris-HCl, pH 7.5 at RT, 0.1 mM EDTA). His_6_ tags were cleaved by adding His_6_-tobacco etch virus (TEV) protease and incubating overnight at 4 °C. Cleaved tags and TEV protease were removed by a second Ni^2+^ affinity column purification. ARNT PAS-B proteins were further purified using a size exclusion column (HiLoad 16/600 Superdex 75 or 200 pg, Cytiva) in the final buffer for later analyses (44.7 mM Tris-HCl, 5.3 mM phosphate, pH 7.0 at RT, 17 mM NaCl, 1 mM DTT). Proteins were concentrated with an Amicon stirred ultrafiltration unit with a 10 kDa filter to 320 – 400 μM. Samples were flash frozen with liquid nitrogen and stored at –80 °C.

### Compound sources

KG-279 (5-Chloro-1,3-benzodioxole) and KG-655 (3,5-bis(trifluoromethyl)phenol) were purchased from Aldrich, while KG-548 (5-[3,5-bis(trifluoromethyl)phenyl]-1H-tetrazole) was purchased from Fluorochem.

### NMR spectroscopy

NMR experiments were performed on 700 or 800 MHz Bruker Avance III cryoprobe-equipped spectrometers, operating at 298.1 K unless otherwise noted. For all experiments with fixed protein concentrations, protein samples at 240 – 280 µM were prepared by adding 5% (v/v) D_2_O, 1% DMSO, and up to 5 mM of ligands (KG-279, KG-548, or KG-655). 200 μl or 500 μl of the prepared samples were added to 3 mm or 5 mm NMR sample tubes, respectively. Standard 1D ^1^H and ^19^F experiments, as well as ^15^N/^1^H and ^13^C/^1^H HSQC experiments, were acquired with 3 mm NMR tubes, while double-filtered HSQC-NOESY and constant time (CT)-^13^C/^1^H HSQC experiments were acquired with 5 mm tubes. Unless specifically mentioned, all ^13^C/^1^H HSQC spectra were collected at 278.1K to maximize the visibility of an upfield-shifted ligand-bound protein peak. All filtered HSQC-NOESY and ^13^C/^1^H HSQC experiments used to assign the NOE correlations were collected at 298.1K, as were ligand-only 1D ^1^H NMR spectra.

Chemical shift assignments of the ARNT PAS-B WT backbone amides and side chains were obtained from previously published work (BMRB entry 6597) (53). Additionally, for mapping the chemical shift perturbation and assigning the bound state, constant-time ^13^C/^1^H HSQC were recorded using ARNT PAS-B WT (280 µM) and the ligands (KG-655 and KG-279) at concentrations of 0, 0.25, 0.5, 1, 2 and 5 mM. ^1^H and ^13^C resonances of ARNT PAS-B interacting with KG-279, KG-548, and KG-655 were assigned using the standard F3-double-filtered NOESY-HSQC 3D experiment in the Bruker pulse sequence library (hsqcgpnowgx33d) (54–57). An NOE mixing time of 110 ms and a recycling delay of 1.5 s were used. Acquisition times for the indirect dimensions were 9.16 ms (t_1_), 13.4 ms (t_2_), and 68.7 ms (t_3_). Finally, binding was also monitored using ^15^N/^1^H HSQC chemical shift perturbation and comparisons of the ligand ^19^F chemical shift and linewidth in the free and bound state. Stereospecific assignments of valine and leucine methyl groups were achieved using an elegant approach developed by Wüthrich and coworkers, exploiting sign differences between the two methyl signals seen in CT-^13^C/^1^H HSQC spectra acquired with a 26.6 ms constant time on a 10% ^13^C-fractionally labeling sample (58, 59). All NMR data were processed with NMRFx Analyst and analyzed using NMRviewJ (60, 61).

The effect of ligand binding on ARNT PAS-B homodimerization was evaluated by diffusion ordered spectroscopy (DOSY) NMR. 100, 280, and 600 μM protein samples were prepared by adding 5% (v/v) D_2_O, 2% DMSO-D_6_ and 5 mM KG-548, 5 mM KG-655, or 2 mM KG-279. 1D ^1^H NMR spectra were obtained at varying gradient strengths (2-98%, 16 points, 512 scans) to evaluate diffusion in the presence of ligand at varying protein concentrations and processed in Bruker TopSpin (Bruker Spectrospin, Fällanden, Switzerland). Diffusion coefficients (D) were obtained by plotting the relative intensities (I), determined by integrating a portion of the aliphatic region (^1^H 1 – 3 ppm), versus the gradient strengths (g) and fitting to the following equation using Grace (https://plasma-gate.weizmann.ac.il/Grace/), where Io is the intensity at the lowest gradient strength, γ is the gyromagnetic ratio of proton (26752 rad/sec×G), is the gradient pulse width (3.6 ms) and Λ1 is the diffusion delay time (99.9 ms) (62):

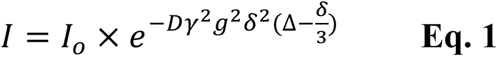

The binding affinities of ARNT PAS-B against KG-548 and KG-655 were estimated using 1D ^19^F NMR. Varying concentrations of ARNT PAS-B (25 µM, 50 µM, 100 µM, 200 µM, 500 µM, and 1 mM) were prepared by adding 5% (v/v) D_2_O, 2% DMSO-D_6_ and 300 μM ligand (KG-548 or KG-655). 1D ^19^F NMR spectra were obtained (4096 scans) at each concentration and processed using Bruker TopSpin and NMRFx (61). Binding affinities were determined by plotting the intensity of the small molecule fluorine peaks (ο_obs_) versus the concentration of ARNT PAS-B ([L]_t_) in Grace. The resulting plot was fit to the following equation to derive binding affinity (K_d_) and maximal peak intensity (ο_max_), where [P]_t_ is the concentration of small molecule (300 μM) (63):

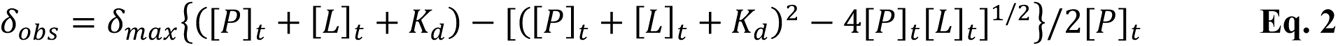

The binding affinity of KG-279 was calculated from the ligand-induced chemical shift changes of selected methyl peaks in a ligand-titration constant time ^13^C-^1^H HSQC experiment. The analysis was performed in NMRviewJ using the Quadratic10 equation (60) and further validated using Grace.

### Weighted ensemble molecular dynamics simulation

To sample the opening of the internal cavity in the ARNT PAS-B domain, we initiated a weighted ensemble molecular dynamics (WEMD) simulation from an equilibrated structure of the receptor in explicit solvent using the WESTPA 2.0 software package (64) and the GPU-accelerated AMBER 22 dynamics engine (65). To prepare this equilibrated structure, we carried out the following steps. Heavy-atom coordinates for ARNT PAS-B were extracted from the crystal structure (PDB code 4EQ1 (48)) and hydrogens were added according to neutral pH using the Reduce tool, as implemented in MolProbity (66). The protein was modeled using the AMBER ff19SB force field (67) and solvated in a truncated octahedral box of OPC water molecules (68) with a 12-Å clearance between the protein and the edge of the box. To ensure a net neutral charge for the simulation system, 5 Na+ and 3 Cl-counterions were added.

Additional ions were included to yield the same salt concentration as our NMR experiments (17 mM NaCl). The solvated system was energy-minimized and then equilibrated in three stages. In the first stage, we performed 20 ps of dynamics in the canonical *NVT* ensemble with a constant number of particles *N*, volume *V*, and temperature *T* of 25 °C, applying positional restraints of 1.0 kcal mol^-1^ Å^-2^ to all heavy atoms of the receptor. In the second stage, we maintained positional restraints on the receptor and equilibrated the system for 1 ns in the isothermal-isobaric *NPT* ensemble with a constant number of particles *N,* pressure *P* of 1 bar, and temperature *T* of 25 °C. In the final stage, we removed all positional restraints and equilibrated the system for 1 ns in the NPT ensemble. Constant temperature and pressure were maintained using a weak Langevin thermostat (frictional coefficient of 1 ps^-1^) and Monte Carlo barostat (pressure changes attempted every 0.2 ps), respectively. To enable a time step of 2 fs, bonds involving hydrogens were restrained to their equilibrium values using the SHAKE algorithm (69). Long-range electrostatic interactions were treated using the particle mesh Ewald method (70) while short-range, non-bonded interactions were truncated at 10 Å.

Our WEMD simulation of ARNT PAS-B was run using a one-dimensional progress coordinate consisting of the solvent-accessible surface area of residues lining the internal cavity of the receptor that contains crystallographic water molecules. These residues were selected based on visual inspection of the internal cavity and consisted of residues F363, S365, F373, I396, R409, S411, F412, V415, V425, F427, F429, M439, T441, S443, I457, C459, and N461. The minimal adaptive binning scheme (71) was applied to the progress coordinate using 15 bins, a target number of 8 trajectories per bin, and a resampling time interval τ of 100 ps for each WE iteration. Our WE simulation was run for 600 iterations (total simulation time of 8 µs) to achieve reasonable convergence of the probability distribution as a function of the progress coordinate.

### Ensemble docking

We determined binding poses of the KG-655 and KG-279 ligands in the internal cavity of the ARNT PAS-B receptor by docking each ligand to an ensemble of receptor conformations with NOE-based distance restraints between the ligand and receptor (**Tables S1, S2**). For the KG-655 and KG-279 ligands, 12 and 28 NOE-based distance restraints, respectively, were generated by qualitative analysis of NOESY peak intensities, using these to establish upper limits of 2.8/3.5/5.0 Å for strong, medium, and weak peaks; lower limits were always set to be 1.8Å as van der Waals contact distance. The ensemble of receptor conformations consisted of 10 conformations from a representative, continuous pathway generated by our WEMD simulation mentioned above with varying extents of solvent-accessible surface area for the internal cavity, ranging from 220 to 290 Å^2^. To generate this ensemble, we first clustered a representative, continuous trajectory from our WEMD simulation using a quality threshold clustering algorithm (72) and then selected the conformation closest to the centroid for each of the resulting clusters, yielding an ensemble of 10 receptor conformations. For the clustering calculation, the similarity metric was the heavy-atom RMSD of the protein from the initial, equilibrated structure with a quality threshold (cluster radius) of 2 Å.

All docking calculations were performed using the HADDOCK software package (73). As implemented in HADDOCK, our docking protocol consisted of three stages: (i) randomization of orientations and rigid-body energy minimization with ambiguous interaction restraints (AIRs) between the ligand and receptor to generate 1000 docked poses for each of the 10 receptor conformations (total of 10,000 models), (ii) semi-flexible simulated annealing in torsion space of the 1000 top-scoring poses, and (iii) refinement of each top-scoring pose using a short molecular dynamics simulation with an 8-Å shell of explicit solvent at 25 °C and position restraints on non-interface atoms.

For the first stage of the docking calculation, ambiguous ligand-receptor restraints were specified for a set of “active” NOE-based methyl-containing residues of the receptor that interact with the ligand. For the KG-655 ligand, seven active residues were specified: I396, L408, V415, L418, L423, V425 and I457. For the KG-279 ligand, the same active residues were specified except for L418. For the subsequent stages of the docking calculation (semi-flexible annealing and water refinement) we specified unambiguous ligand-receptor restraints as a set of inter-heavy-atom distances that were obtained by extending each NOE-derived interproton distance by 1 Å to implicitly account for the two C-H bonds. For the KG-655 and KG-279 ligands, this set consisted of 19 (12 involving non-equivalent heavy atoms) and 31 distances (28 involving non-equivalent heavy atoms), respectively. Parameters for the receptor and ligand were taken from the OPLS united-atom force field (74) and default HADDOCK settings for protein-ligand docking were applied. For each ligand, the resulting 1000 top-scoring poses from the docking calculation were clustered using the RMSD of the ligand from the best-ranked pose after alignment on the Cα atoms of the receptor and a clustering cutoff of 1.5 Å. To avoid double-counting certain docked poses due to the two-fold symmetry of the KG-655 ligand, the ligand RMSD involved only three heavy atoms that lie along the symmetry axis: two carbon atoms of the aromatic ring and the oxygen on the hydroxyl substituent of the ring. For the KG-279 ligand, the ligand RMSD involved all heavy atoms of the ligand. The resulting nine clusters were then ranked by the average HADDOCK score of the top four poses of each cluster. We selected the top-scoring pose of each cluster; for the first cluster, the pose with the second-best score was selected due to the best-scoring pose having unfavorable electrostatic interactions. We also discarded models with NOE distance violations >0.3 Å. These two filters reduced the 1000 docked poses for each ligand to a final set of 5 poses.

For each ligand, we further refined the final five poses with all-atom models by adding in nonpolar hydrogens and running a 21-ns conventional molecular dynamics (cMD) simulation for each pose using the AMBER 20 dynamics engine (65). All cMD simulations were run in the NPT ensemble (25°C and 1 atm) with the AMBER ff19SB force field (67) for the ARNT PAS-B receptor, GAFF2 (75) parameters for the small-molecule ligands, OPC model for the explicit water molecules (68), Joung-Cheatham parameters for neutralizing Na^+^ and Cl^-^ ions (as created for the TIP4P-EW model) (76), and NOE-based interproton distance restraints between the ligand and receptor using square-bottom wells that account for the corresponding lower and upper bounds. Distance restraints were gradually applied during the first 10 ns by increasing the force constant from 0.2 to 2 kcal mol^-1^ Å^-2^; a force constant of 2 kcal mol^-1^ Å^-2^ was then applied throughout the latter 10 ns. Finally, the force constant was gradually reduced to zero during a 1 ns simulation while applying positional restraints of 1.0 kcal mol^-1^ Å^-2^ to all receptor heavy atoms except for the active sidechains. GAFF2 ligand parameters (**Tables S3, S4**) were derived by fitting partial atomic charges to electrostatic potentials using the restricted electrostatic potential (REsP) method (77); electrostatic potentials were calculated at the HF/6-31G* level of theory using the Gaussian 16 A03 software package.

Prior to running the above cMD simulation, the ligand-receptor system was prepared by immersing the system in a truncated octahedral box of water molecules with a 12 Å clearance between the system and the edge of the box. To ensure a net neutral charge, 5 Na^+^ and 3 Cl^-^ counterions were added. Additional ions were included to yield the same salt concentration as our NMR experiments (17 mM NaCl). The solvated system was energy-minimized and equilibrated using the same protocol described above for WE simulations of the ligand-free receptor with the exception of applying the following during all three stages of equilibration: positional restraints of 1 kcal mol^-1^ Å^-2^ to all heavy atoms of the receptor except for the active sidechains involved in NOE-based interproton distance restraints and interproton distance restraints with a force constant of 0.2 kcal mol^-1^ Å^-2^.

## Results

### KG-548 binds to the surface of ARNT PAS-B and promotes homodimerization

We recently proposed KG-548 as a surface-binding ligand based on high-pressure NMR data, contrary to our prior assumption that this ligand binds within the 105 Å^3^ of interconnected internal cavities (49). This proposal was supported by the crystal structure of a ARNT PAS-B homodimer complexed with KG-548, with a single molecule of the ligand bound near the dimerization interface generated by the β-sheet surfaces of the two ARNT PAS-B molecules (49). While consistent with the high-pressure NMR data, the ARNT PAS-B domain is not known to homodimerize outside of high concentration solutions (53) or crystallization settings. In addition, the same site has been shown to interact with crystallization reagent polyethylene glycol (PEG) in a previously solved X-ray structure of an ARNT PAS-B homodimer (48). These observations raise questions of whether KG-548 binds to the same surface site in solution.

First, we investigated the impact of KG-548 on ARNT PAS-B homodimerization in solution using diffusion ordered spectroscopy (DOSY) experiments. DOSY is an NMR method that reports on the diffusion coefficients (D) of molecules in the sample, which are inversely proportional to the hydrodynamic radius of the molecules (62, 78). If ARNT PAS-B equilibrium shifts towards dimer from monomer, the observed diffusion coefficient is expected to become smaller. Our initial step involved measuring the diffusion coefficient of ARNT PAS-B at three concentrations (100, 280, 600 µM). By assuming a 100% monomeric state at the lowest concentration, we extrapolated the monomeric proportions at higher concentrations (**Table S5**). This analysis indicated a concentration-dependent trend towards dimerization for ARNT PAS-B, as expected (53). From this foundation, we then evaluated the monomer/dimer equilibrium of ARNT PAS-B in the presence of 5 mM KG-548, finding that this ligand indeed promotes homodimerization, most notably at low protein concentration. To validate that the observed differences were not attributable to changes in sample viscosity, we verified that the diffusion coefficient of water remained constant at all protein and ligand concentrations used here.

Next, we explored the importance of surface residues located near the crystallographically-observed binding site using both protein– and ligand-observed solution NMR methods (49). To examine binding from the perspective of the protein, we used solution ^13^C/^1^H HSQC experiments to map the effects of adding KG-548 to ARNT PAS-B, observing ligand-induced chemical shift changes in several sites (**Fig. 2A**); we focused our analyses here on Ile δ1 methyl groups given their distribution throughout the ARNT PAS-B domain and suitable NMR properties (upfield ^13^C shifts, narrow linewidths). We complemented this with ligand-observed 1D ^19^F NMR experiments, taking advantage of the two trifluoromethyl groups in KG-548 (**Fig. 2A**). Comparing ^19^F spectra acquired on KG-548 alone and in the presence of ARNT PAS-B, we saw the appearance of a broad, protein-dependent peak at ∼ –61.5 ppm, which we designate as the bound peak of the ligand (**Fig. S1**). This peak showed temperature-dependent line narrowing, consistent with fast exchange kinetics at 298.1K (**Fig. S1**). In addition, we measured a K_D_ ∼ 350 μM binding affinity for the ligand using ^19^F NMR, by titrating varying concentrations of ARNT PAS-B (25 – 1000 μM) into 300 μM KG-548 (**Fig. S2**).

**Fig. 2.**
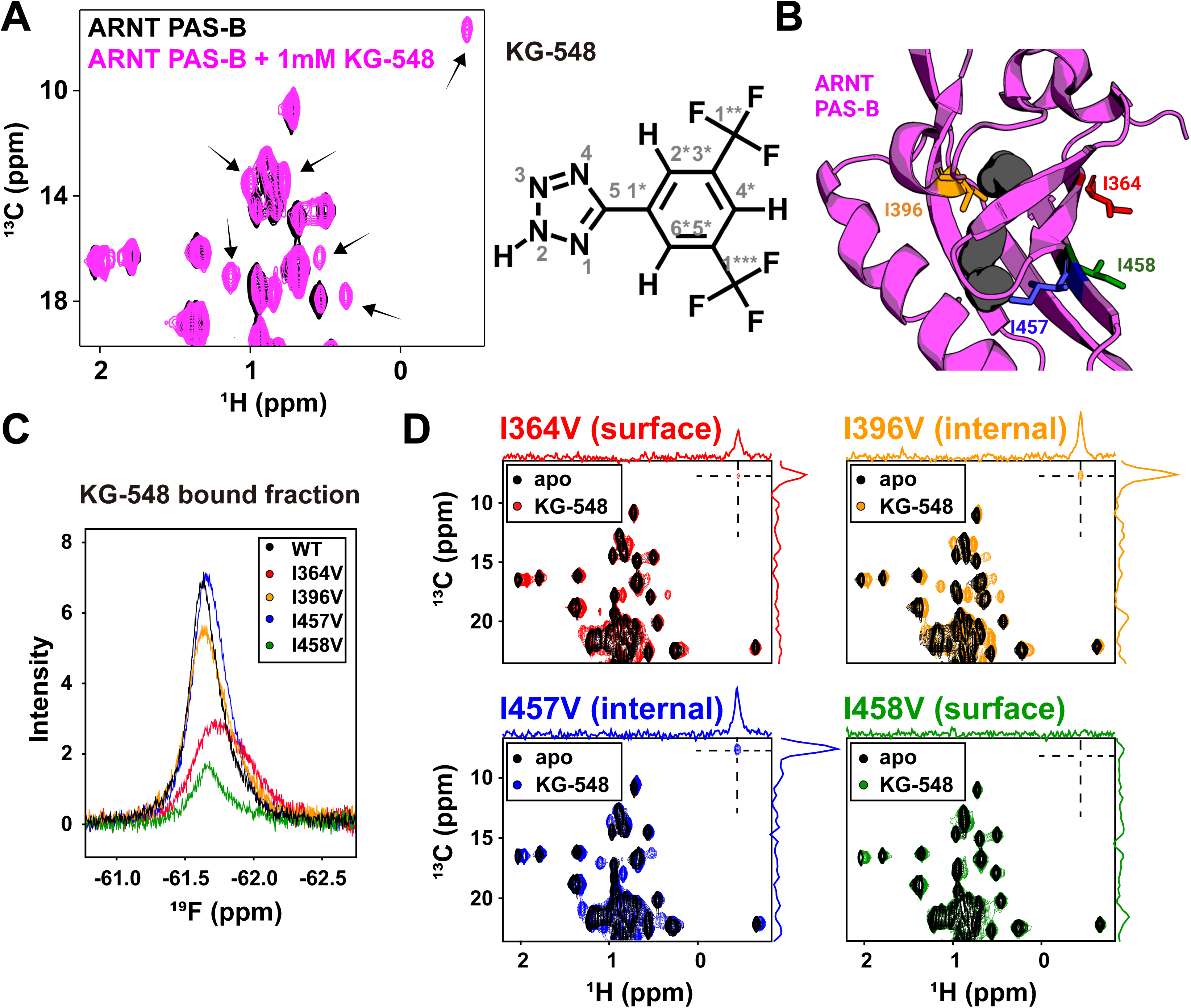
Isoleucine residues I364 and I458 are important to the binding of KG-548. A) ^13^C/^1^H HSQC spectra of 250 μM protein without and with 1 mM of KG-548. Several methyl peaks (black arrows), and in particular, peaks in the Ile region, showed chemical shift perturbations in the presence of the ligand. B) Four residues were selected for Ile to Val mutagenesis studies: two surface-facing Iles (I364 and I458) and two cavity-facing (I396 and I457). C) ^19^F NMR spectra of 1 mM KG-548 incubated with the 250 μM of ARNT PAS-B mutants. Overlaid spectra are zoomed in on the bound fraction of the ligand. Mutations I364V and I458V substantially reduced binding. D) ^13^C/^1^H HSQC spectra of 1 mM KG-548 added to 250 μM ARNT PAS-B mutants. Disruptions of binding were observed for the I364V and I458V mutants, as particularly clear by the reduction/disappearance of the upfieldshifted ligand-dependent Ile δ1-methyl peak.

From this foundation, we investigated the contributions to binding of four isoleucine residues with separate Ile to Val point mutations, including two surface Iles located at the KG-548 binding site (I364, I458) as well as two internal ones (I396 and I457) that face the previously identified internal cavities (**Fig. 2B**) (48). From both ligand– and protein-detected approaches, we observed that the I364V and I458V point mutations markedly reduced KG-548 binding affinity, as seen by both proteins showing lower bound fractions of KG-548 in 1D ^19^F NMR (**Fig. 2C**) or in the number and magnitude of ligand-induced Ile 81 ^13^C/^1^H HSQC peak shifts (**Fig. 2D**). This reduction in binding was particularly stark for I458V, with minimal chemical shift changes observed upon the addition of the ligand, and a total loss of an upfield-shifted Ile-methyl peak (**Fig. 2D**). These data and prior observations from the crystal structure led us to conclude that this ligand-dependent upfield-shifted peak is the δ1-methyl of I458; in contrast, minimal perturbation to protein/ligand interactions were seen for the two mutants of the cavity-facing I396 and I457. Importantly, we confirmed that none of the four point mutations perturbed the overall fold of ARNT PAS-B as judged by comparisons of mutant and wildtype ^15^N/^1^H HSQC and ^13^C/^1^H HSQC spectra (**Figs. S3, S4**). Taken together, these solution data demonstrate the importance of the I364 and I458 side chains in binding KG-548, in strong agreement with information provided by the crystal structure: these surface-orientated Ile residues are essential for binding.

To determine the solution binding pose of KG-548 on the ARNT PAS-B surface independent of the crystal structure, we acquired ^13^C, ^15^N double-filtered ^13^C/^1^H HSQC-NOESY data to selectively measure intermolecular NOE correlations between the ligand and the protein methyl groups (54–57). To guarantee a high concentration of the bound state, we added a maximum soluble concentration (5 mM) of KG-548 to uniformly ^13^C/^15^N-labeled ARNT PAS-B (280 µM), optimizing our ability to measure intermolecular NOEs between protein and ligand protons within ∼ 5 Å of each other. KG-548 has three hydrogens, including two chemically equivalent ones, resulting in two separate peaks in the 1D ^1^H spectrum that we could unambiguously assign (**Fig. 3A**). NOEs to either of these ligand signals were correlated to protein ^13^C chemical shifts that we assigned by the chemical shifts seen in a ^13^C/^1^H HSQC spectrum of the same sample (**Fig. 3A**), which in turn were referenced against previously-deposited side chain chemical shift assignments from our group (53) and confirmed by mutagenesis studies as described above (**Fig. S4**). As expected, these NOESY data show that I364 and I458 are both near the ligand binding site. Furthermore, strong NOE correlations were observed between the non-degenerate proton of the ligand and methyl groups of I364 (γ2) and I458 (81, γ2), and a weaker NOE correlation was observed between the degenerate protons and the I458 C81 position (**Fig. 3A**). Notably, the crystal structure agrees with these distances, again confirming the validity of the surface binding site we previously identified (**Fig. 3B**).

**Fig. 3.**
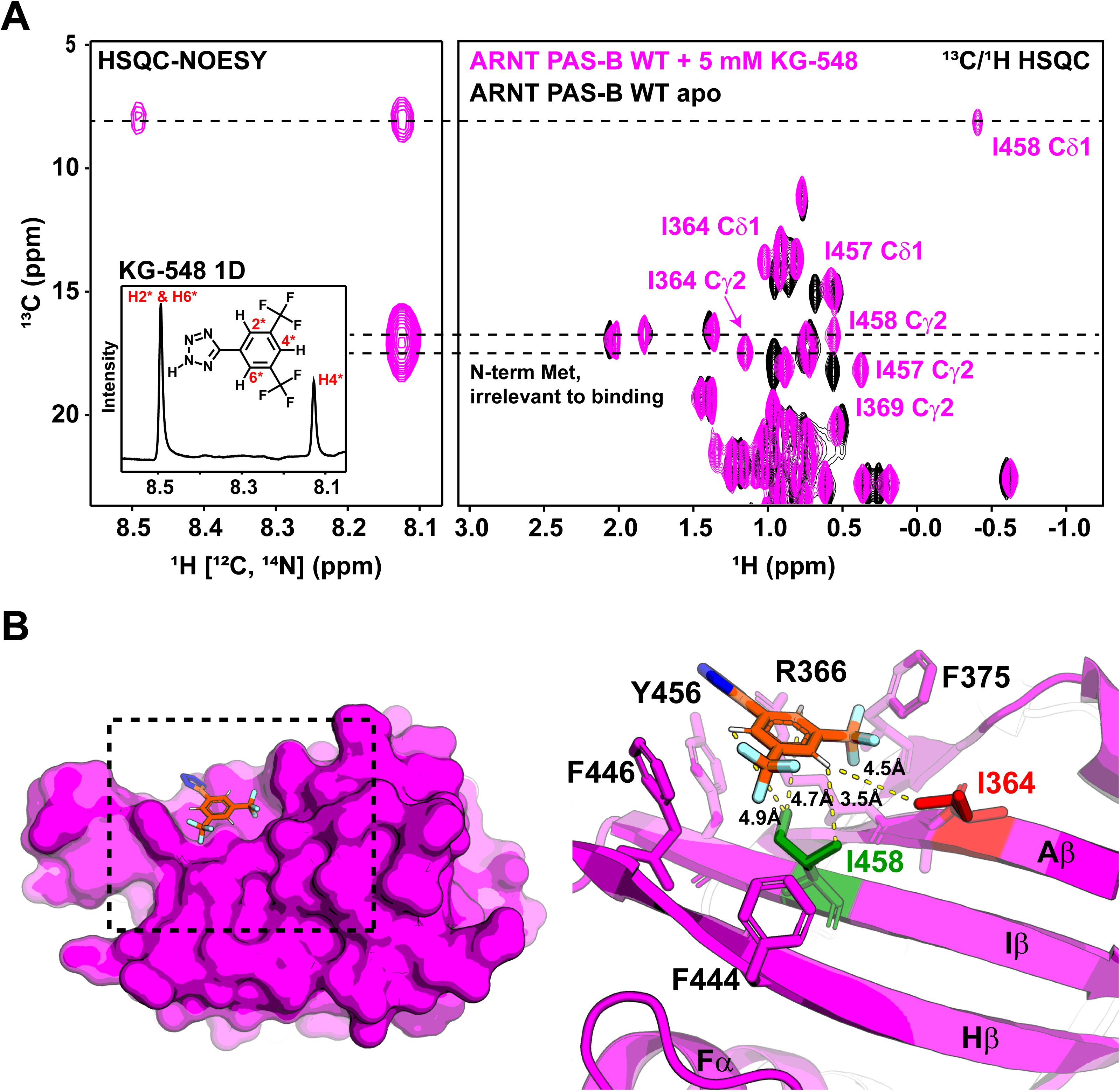
KG-548 binds to the β-sheet surface of ARNT PAS-B. A) ^13^C, ^15^N double filtered ^13^C/^1^H HSQC-NOESY experiment shows NOE correlations between KG-548 and surface Iles I364 and I458. Assignments were made by aligning the HSQC-NOESY spectrum with the ^13^C/^1^H-HSQC spectrum of ARNT PAS-B with KG-548. 1D ^1^H NMR of KG-548 is shown in the inset on the left. B) Diagrams of the ARNT PAS-B crystal structure in complex with KG-548 (PDB: 8G4A). Residues within 5Å of the ligand are highlighted with sticks (right). Distances between protons on KG-548 and surrounding methyl groups are shown (yellow dashed lines).

To investigate whether we could completely disrupt the surface binding site of KG-548 by removing both residues I364 and I458, we purified the ARNT PAS-B I364V/I458V double mutant and performed the same 1D ^19^F NMR experiments. As shown in **Fig. S5**, this mutant further reduced binding compared to either single point mutation but was insufficient to remove binding entirely. Based on the crystal structure reported earlier, several aromatic residues (F375, F444, F446, Y456) were also found in proximity to the binding site of KG-548 (**Fig. 3B**), one of which (Y456) is substantially displaced compared to the apo ARNT PAS-B structure. We suspect that one or more of these residues may contribute hydrophobic interactions involved in keeping the ligand in place.

### KG-655 exhibits dual binding modes

KG-655 is a structural fragment of KG-548 with a hydroxyl replacing the tetrazole group of the latter compound (**Fig. 4A**). Despite the structural similarities of these compounds, our prior high-pressure NMR data suggested a different binding mode between them, implicating that KG-655 binding (and not KG-548) could reduce the void volume of ARNT PAS-B, consistent with it being an internal binder (49). Therefore, we set out to test this hypothesis using the aforementioned approaches. First, we found that KG-655 also promotes ARNT PAS-B homodimerization, but to a lesser extent than KG-548, from DOSY experiments (**Table S5**). We also collected ^13^C/^1^H HSQC spectra of the ARNT PAS-B domain in the presence of KG-655 (**Fig. 4A**). At first glance, the same isoleucine residues were perturbed, including the upfield-shifted I458 δ1 methyl group, suggesting KG-655 shares the same surface binding interface as KG-548. We then collected ^13^C/^1^H HSQC spectra of the four ARNT PAS-B mutants in the presence of 1 mM KG-655, again demonstrating that the surface I364V and I458V mutations substantially perturbed compound binding, as particularly indicated by the reduction in peak intensity of the I458 δ1 methyl group (**Fig. 4B**). Interestingly, while the I458V mutant showed almost no chemical shift perturbations upon adding KG-548, several methyl groups showed clear chemical shift changes when KG-655 was added, all of which were on inward-facing side chains adjacent to the internal cavity based on our prior ARNT PAS-B assignments (53). Notably, these chemical shift perturbations were also seen in the WT protein and the other three mutants in the presence of KG-655 but not when KG-548 was added (**Fig. 4B**), strongly implicating that KG-655 can access a second, distinct binding mode. Additional support for this hypothesis was provided by ^15^N/^1^H HSQC spectra of the surface-binding disrupting ARNT PAS-B I458V mutant in the absence and presence of 5 mM KG-655 (**Fig. 4C**). Again, several peaks showed broadening and/or chemical shift perturbations compared to the apoprotein, including many perturbed residues that are not located on the β-sheet surface, consistent with a second binding mode for KG-655.

**Fig. 4.**
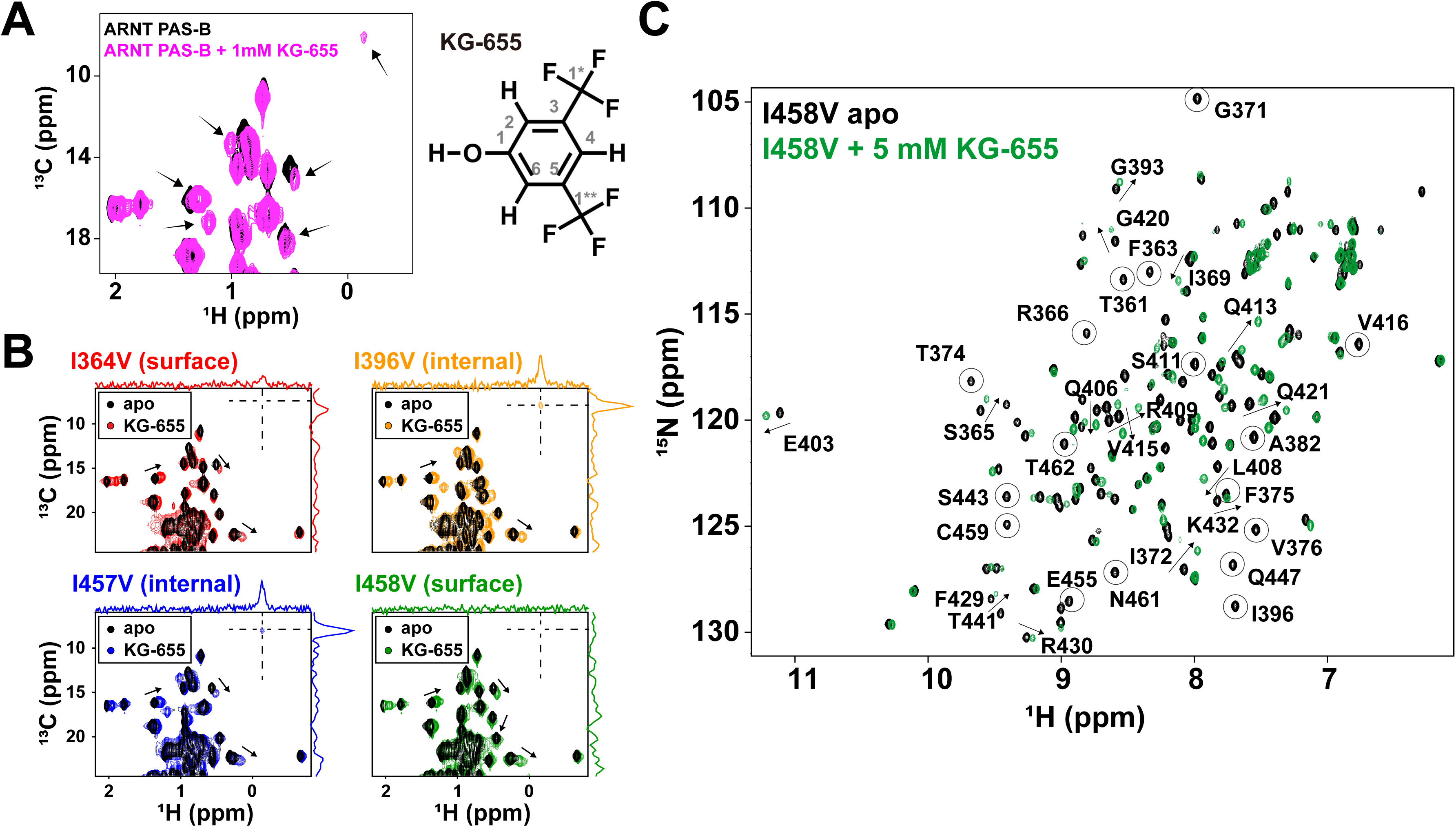
KG-655 exhibits two distinct binding modes. A) ^13^C/^1^H HSQC spectra of 250 μM with and without 1 mM of KG-655. Residues showing chemical shift perturbations are marked with black arrows. B) ^13^C/^1^H HSQC spectra of 1 mM KG-655 added to 250 μM ARNT PAS-B mutants. Mutations I364V and I458V disrupted KG-655 binding, similar to their effects on KG-548. Additional chemical shift perturbations were observed independent of the surface Ile mutations (black arrows). C) ^15^N/^1^H HSQC spectra of ARNT PAS-B I458V with and without 5 mM of KG-655. Intermediate to fast exchange behavior was observed, as indicated by the broadening (black circles) and/or chemical shift changes (black arrows) of apo cross-peaks in the presence of the ligand. Many perturbed peaks are not located on the β−sheet surface (e.g., I396, E403, L408, G420).

Evidence of a dual binding mode for KG-655 was also seen in ligand-observed ^19^F NMR (**Fig. S1**), which, like KG-548, exhibited a protein-bound peak for KG-655 when mixed with ARNT PAS-B (∼ –61.5 ppm), downfield-shifted from the free-ligand peak. Given the similarity of the chemical shift for this peak between KG-548 and KG-655, we again interpret this as representing the surface-bound fraction of the ligand. However, in addition to this peak, the free-ligand peak was also broadened, which was not seen for KG-548. We speculate that this broadening resulted from the second binding mode. We again titrated ARNT PAS-B into 5 mM KG-655 in an attempt to estimate the binding affinities (**Fig. S2**); we estimate millimolar affinity binding of this compound, but the low affinity of this interaction and dual binding modes of KG-655 complicate rigorous quantitation.

To determine the precise binding site(s) of KG-655, we again collected the ^13^C, ^15^N double filtered ^13^C/^1^H HSQC-NOESY experiments using wildtype ARNT PAS-B. Compared to KG-548, we saw substantially more intermolecular NOE correlations between the KG-655 and the WT protein (**Figs. 5A, B, C**). To help with assignments, we aligned the constant time ^13^C/^1^H HSQC spectrum of ARNT PAS-B with 5 mM KG-655 with the HSQC-NOESY experiments showing both H-C and H-H dimensions (**Fig. 5A**). Notably, all Leu and Val methyl groups were stereospecifically assigned using constant time ^13^C/^1^H HSQC spectra recorded on a 10% randomly U-^13^C labeled sample, using different signs of peaks in these spectra to distinguish pro-R and pro-S methyl groups (58, 59) (**Fig. S6**). In this way, we assigned all methyl peaks showing NOE correlations with KG-655, including several residues proximate to the internal cavity (I396, L408, V415, L418, L423, V425, I457), a residue on the surface (I458), and an outlier (V381) that is distant from both binding sites. This data unambiguously placed the second binding mode of the KG-655 inside the ARNT PAS-B cavity.

**Fig. 5.**
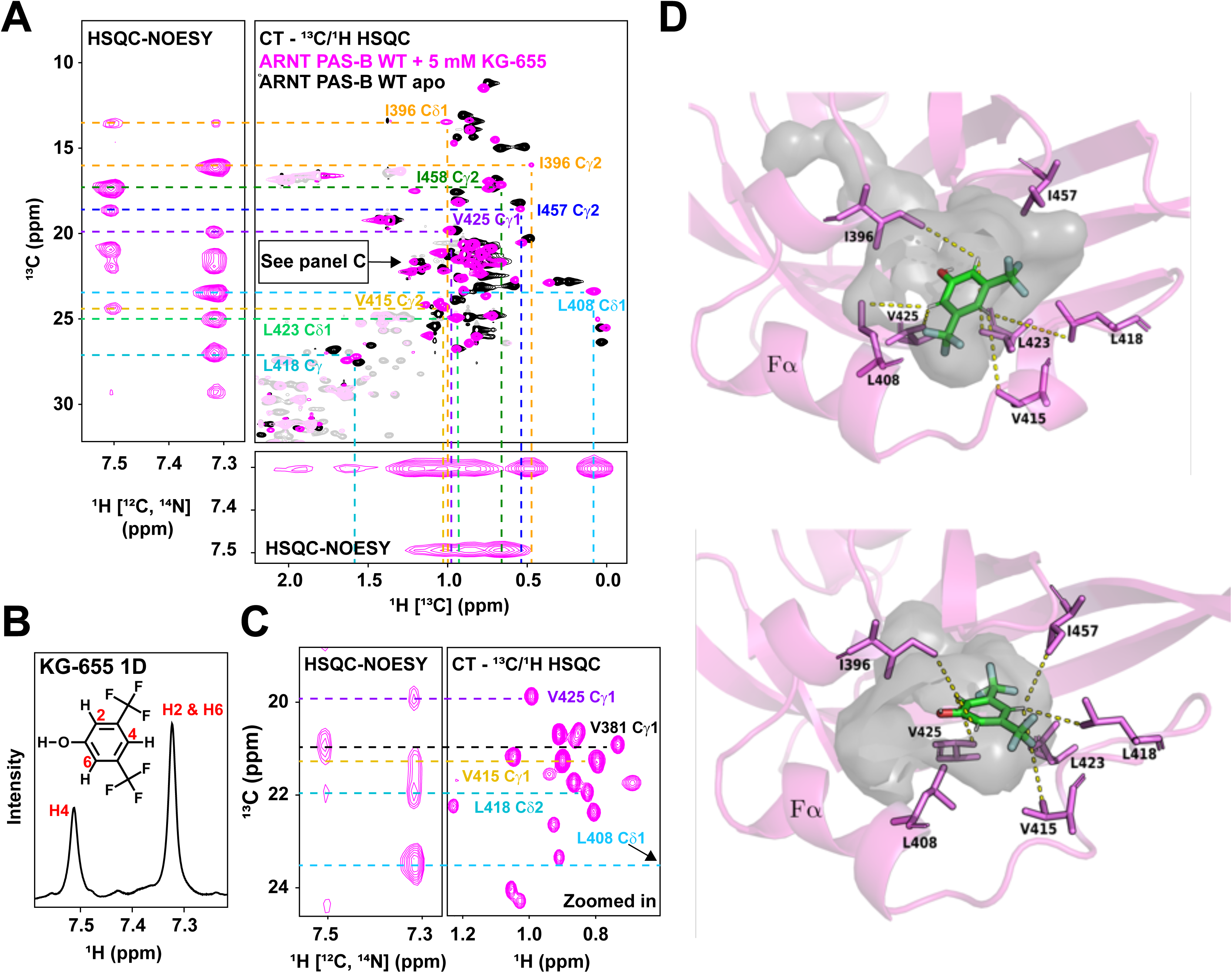
KG-655 binds to both the surface and internal cavities of ARNT PAS-B. A) ^13^C, ^15^N, double filtered ^13^C/^1^H HSQC-NOESY experiment revealed residues that are close to KG-655 binding site(s), which included cavity-facing residues (I396, L408, V415, L418, L423, V425, I457) and a surface-facing residue (I458), with one outlier distant from both sites (V381). NOE correlations are assigned by aligning the HSQC-NOESY spectra with a constant time (CT)-^13^C/^1^H HSQC spectrum of ARNT PAS-B with 5 mM KG-655. B) Zoomed-in view of part of panel A. C) 1D ^1^H NMR of KG-655. D) Two potential binding poses of KG-655 (green) in the hydrated internal cavity (gray) of ARNT PAS-B (magenta) from ensemble docking. Among the seven receptor residues (magenta sticks) involved in NOE-based, ligand-receptor distance restraints, six (top pose) and five residues (bottom pose) have satisfied restraints. For each of these residues, the shortest satisfied distance is indicated by a dashed yellow line between the methyl carbon and ligand proton.

We compiled all the observed NOE correlations that are consistent with the cavity binding mode (listed in **Table S1**) and performed ensemble docking to visualize the proposed binding mode of KG-655 inside the cavity. To generate ARNT PAS-B conformations with sufficiently open internal cavities for accommodating the ligand, we applied the weighted ensemble simulation strategy (79, 80), which enhances the sampling of rare events (e.g., large-scale conformational transitions in proteins (81), binding processes of proteins (82, 83) and DNA (84)) without introducing any bias into the dynamics. As an ensemble, the five top-scoring poses satisfied 10 of the 12 interproton distance restraints between the ligand and receptor involving non-equivalent protons. The two unsatisfied distance restraints involved the only two receptor residues for which both methyl groups participate in distance restraints (I396 and L418). In these cases, restraints were satisfied for one methyl group but not the other, potentially due to artifacts by spin diffusion. **Fig. 5D** shows the two poses with the highest number of satisfied distance restraints, which involved either six or five of the seven receptor residues involved in the observed NOE correlations. Given the NOE data, we strongly believe the ligand adopts multiple orientations within the cavity to satisfy all distance restraints.

Finally, to assess whether the two binding modes are related, we acquired the same double-filtered HSQC-NOESY experiment using the ARNT PAS-B I364V/I458V double mutant, which should severely disrupt surface binding (**Fig. S7**). We concluded that the NOE correlations from the internal-facing residues were unaffected by assigning all methyl groups close to the ligand. As expected, the correlation between the ligand and surface residue I458 was the only one that disappeared. This data suggests that the disruption of surface binding did not influence internal binding.

### Conformation-destabilizing mutant Y456T completely disrupts KG-655 surface binding

Since the two binding modes of KG-655 are independent, we asked whether we could remove surface binding altogether to study the TACC3-disrupting effects of the internal binding mode alone. As shown above, individual or paired mutations to the key I364 and I458 residues were insufficient to eliminate surface binding completely. We turned our attention to residue Y456, another residue that showed a marked conformational change in the crystal structure of the ARNT PAS-B/KG-548 complex (**Figs. 3B, S8**). We previously reported that mutations to Y456, along with other aromatic residues in the surrounding, could destabilize the wildtype conformation of ARNT PAS-B. Y456T, for example, slowly interconverts between two stably folded conformations: a wildtype (WT) conformation and an alternative “SLIP” conformation, named for a 3-residue shift in the β-sheet register (85, 86). We demonstrated that both KG-548 and KG-655 specifically bound the WT conformation, allowing them be used to modulate SLIP/WT equilibrium (87). Here, we monitored the surface-bound fraction of KG-548 and KG-655 mixed with ARNT PAS-B Y456T mutant using 1D ^19^F NMR (**Fig. S9**). While the binding of KG-548 was only slightly perturbed by the mutation, the surface binding of KG-655 was completely disrupted. This is evidenced by the complete loss of the surface-bound peak of KG-655 in the ^19^F spectrum.

In contrast, KG-655 could still bind to the internal site within the Y456T variant, albeit to a lesser extent than its interactions with the WT protein (**Fig. S9**). We have previously estimated the apparent K_d_ of KG-655 to ARNT PAS-B Y456T to be in the range of high μM to low mM (87). Since KG-655 does not bind to the surface of ARNT PAS-B Y456T, this estimation could be interpreted as the binding affinity of the internal binding mode alone. Furthermore, KG-655 does not contain a tetrazole group, which in KG-548 forms polar contacts with R366 on the ARNT PAS-B β-sheet surface for heightened stability (**Figs. 3B, S8**). Therefore, we hypothesize that the smaller KG-655 binds less tightly to the surface site and relies heavily on hydrophobic interactions with surrounding aromatic residues (**Fig. 3B**). Consequently, mutating Y456 was more detrimental to KG-655 surface binding than KG-548.

### KG-279 binds to the internal cavity of ARNT PAS-B

In addition to KG548 and KG-655, our lab previously identified several other low affinity ARNT PAS-B binding ligands using an NMR-based fragment screen (48). We pondered whether other ligands bound ARNT PAS-B in a similar fashion to KG-548 and KG-655. Here, we present data from one such ligand (KG-279, **Fig. 6A**), which showed the most ligand-induced chemical shift perturbations in ARNT PAS-B with ^15^N/^1^H– and ^13^C/^1^H-based titration experiments. From the chemical shift changes of 6 methyl peaks, we estimated around 1.2 mM binding affinity (**Fig. S10**). While KG-279 is not known to disrupt ARNT PAS-B/TACC3 interactions, our preliminary analysis suggests it is also an internal binder (**Fig. 6B**). This ligand contains five peaks in 1D ^1^H NMR (**Fig. 6A**), allowing more distance restraints to be generated using the double-filtered HSQC-NOESY experiment. As with KG-655, we compiled all the NOE correlations for KG-279 (**Table S2**). We found that most correlations were from methyl-containing residues near the internal cavity, with only a weak correlation between the ligand and I458 Cγ2. The titration experiment with up to 5 mM of KG-279 also did not yield the signature upfield shifted peak (I458 Cδ1) in ARNT PAS-B that represents surface binding. Taken together, we concluded that KG-279 is primarily an internal cavity binder, differing from both KG-548 (surface binding only) and KG-655 (both surface and internal binding). **Fig. 6C** shows the top two poses from ensemble docking with the highest number of satisfied distance restraints, which involved five of the six residues observed in the NOE correlations. Like KG-655, docking suggests KG-279 adopts several orientations within the internal cavity of ARNT PAS-B. Lastly, we also tested whether this internally bound ligand would promote ARNT PAS-B homodimerization. Initial observations indicated that at a concentration of 2 mM, the ligand significantly impacted the sample’s viscosity. After compensating for this variation, our data revealed that KG-279 also promotes homodimerization, but had the least influence compared to that of the other two ligands (**Table S5**), underscoring the differential impact of ligands with different binding modes on the dimerization process.

**Fig. 6.**
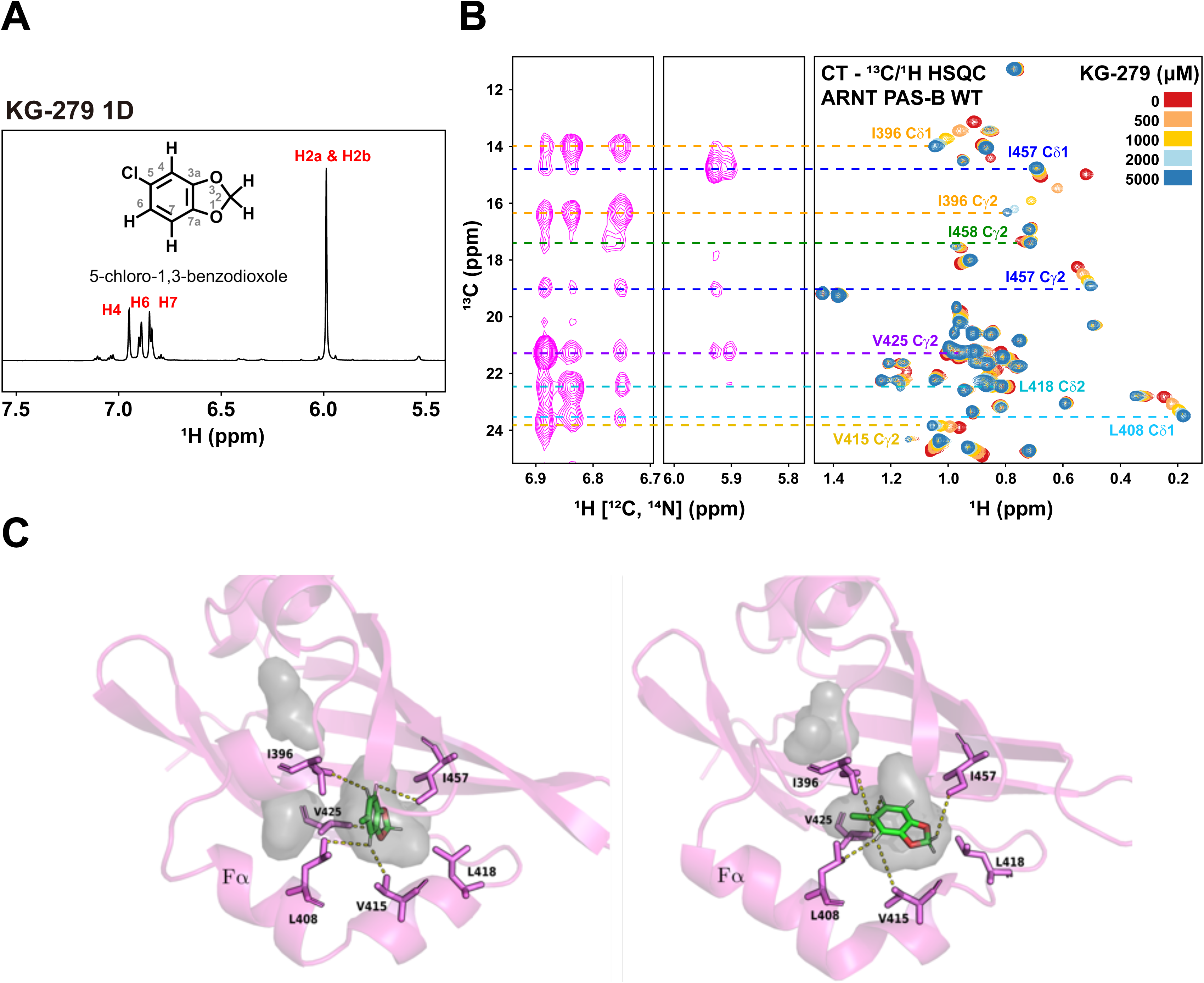
KG-279 shares the same internal binding site as KG-655. A) 1D ^1^H NMR of KG-279. B) Double-filtered HSQC-NOESY experiment showed that the identical cavity-facing residues are also in proximity to bound KG-279. The right shows the constant time (CT)-^13^C/^1^H HSQC spectra of a KG-279 titration (0 to 5 mM) into 240 μM ARNT PAS-B. The titration experiment was also used to approximate the binding affinity of KG-279, which was 1.19 +/− 0.07 mM. Residues showing NOE correlations were assigned by aligning the HSQCNOESY spectrum with the HSQC spectrum of ARNT PAS-B in the presence of 5 mM KG-279. C) Two potential binding poses of KG-279 (green) by ensemble docking in the internal cavity (gray) of ARNT PAS-B (magenta) from ensemble docking. Among the six receptor residues (magenta sticks) involved in NOE-based, ligand-receptor distance restraints, five residues have satisfied restraints. The shortest satisfied distance for each of these residues is indicated by a dashed yellow line between the methyl carbon and ligand proton.

## Discussion

Targeting ARNT/coactivator interactions represents a new opportunity to artificially regulate bHLH-PAS associated signaling pathways (88). Our lab has previously studied several such ARNT-interacting transcriptional coactivators, known as the coiled-coil coactivators (CCCs), named after their shared use of homodimeric coiled coils at their C-terminal ends (41, 43–47). Mechanistically, CCCs are likely recruited to AHR and all three HIF isoform complexes via the ARNT PAS-B domain (41, 45, 46). Consequently, disrupting ARNT/CCC interactions with small molecule inhibitors may represent an effective strategy to achieve inhibition across multiple HIF complexes. This approach also offers a novel approach to modulate the AHR-mediated xenobiotic pathway without perturbing the ligand-recognition of the AHR subunit.

As the first step in this aim, it is critical to understand how small molecules interact with ARNT PAS-B. Most of our ARNT PAS-B binding small molecules previously identified have only moderate binding affinities (48), complicating the binding site(s) characterizations. Here, we couple NMR chemical shift changes, NOESY distance measurements, and ensemble docking, improving our understanding of how three small-molecule ligands differently interact with ARNT PAS-B: KG-548 directly interferes ARNT/coactivator association by competing with the coactivator on the β-sheet surface; KG-655 shares the same surface binding site as KG-548, but also binds to the internal cavity of ARNT PAS-B; KG-279 has not been previously identified as a coactivator inhibitor, but it binds predominately to the water-accessible internal cavity, analogous to where other PAS domains often bind natural or artificial small molecule ligands (35, 89).

Surface-binding ligands are rare in PAS domains, with proflavine binding to HIF-α PAS-B:ARNT PAS-A interface of the HIF bHLH-PAS heterodimer being one of the few documented cases (90–92). Ligands bound to the surface of ARNT PAS-B have not been reported in the past, and indeed, when we first discovered KG-548 as an ARNT PAS-B binding ligand, we assumed it was going into the internal cavities interpreting from ^15^N/^1^H HSQC-based titrations (48).

However, high-pressure NMR data recently suggested that KG-548 may be a surface-binder instead, as the pattern of pressure-dependent chemical shift changes of KG-548-bound ARNT PAS-B differed from that of a protein with an internally bound ligand (49). We subsequently solved an ARNT PAS-B:KG-548 cocrystal structure, showing additional support that the ligand was located on the surface (49). Here, we investigated the binding process in solution using a combination of NMR experiments, including most notably detecting the NOE correlations between the ligand and the protein. We confirmed KG-548 as a surface binder and demonstrated that the ligand promotes ARNT PAS-B homodimerization, removing doubts that the binding to the shallow surface pocket was an artifact of the crystal packing. As noted above, one of the CCCs, TACC3, is likely recruited to the same β-sheet surface that KG-548 binds to, suggesting the ligand would directly compete with the transcriptional coactivator and interfere with ARNT/TACC3 formation (44). Interestingly, this surface has also been shown to interact directly with the HIF-2α PAS-B domain during HIF heterodimerization (**Fig. S11**) (92). Hence, ARNT PAS-B is playing a complex role here, potentially mediating the recruitment of multiple protein binding partners. The mechanistic details of how this is accomplished are beyond the scope of this manuscript but certainly warrant further investigation.

It is unsurprising that KG-655, a structural fragment of KG-548, would share the same surface binding mode with the larger compound. However, interestingly, we found that KG-655 also binds to an internal cavity of ARNT PAS-B, independent of surface binding. The smaller size of KG-655 is likely responsible for this internal binding mode, as KG-548’s tetrazole group was replaced with a hydroxyl group (**Figs. 2A, 4A**). The apo form of ARNT PAS-B contains two adjacent cavities, a larger 65Å^3^ water-containing cavity and a 40Å^3^ secondary cavity. While these cavities are small, their locations are analogous to the larger 300Å^3^ cavity inside the HIF-2α PAS-B domain (35), which is the binding site of the FDA-approved HIF-2α inhibitor, belzutifan (37, 93). Therefore, an ARNT PAS-B internal cavity binder could theoretically block ARNT/coactivator association through a similar mechanism, complementing the direct competition possibly by something binding externally on the ARNT PAS-B β-sheet surface. To our knowledge, this is the first time that direct evidence has been reported showing a ligand could enter the ARNT PAS-B internal cavities. We believe this is a starting point for developing tool compounds and leads for therapeutic strategies that complement the previously described ligands targeting the HIF-α counterparts.

We attempted to test whether we could isolate the internal binding mode for KG-655 by eliminating the surface binding of the ligand. Initially, we removed the two critical surface isoleucine residues, which was insufficient to abolish surface binding entirely (**Fig. S5**). Surprisingly, we found that a conformation destabilizing mutant Y456T could disrupt the surface binding of KG-655, but not KG-548. The Y456T mutation enables ARNT PAS-B to adopt two slowly interconverting confirmations, including a WT state that resembles the wildtype protein (85). KG-655 could not bind to the surface of this mutant, including the WT fraction. The mutation, however, had little effect on KG-548, suggesting the surface binding of KG-655 was less stable, which correlates with KG-548 being the more efficacious inhibitor. This observation, coupled with the fact that internal-binding ligand KG-279 does not inhibit ARNT PAS-B/TACC3 association, strongly suggests that the surface binding mode is responsible for the inhibitory effects of KG-548 and KG-655. However, we must note that testing these ligands’ effects on TACC3 binding affinities is a complex endeavor due to these compounds’ low potency and solubility. Further studies will be required to separately evaluate the two binding modes reported and determine their specific role in inhibition.

To visualize the proposed binding mode of KG-655 in the internal cavity of ARNT PAS-B, we performed molecular docking to an ensemble of ARNT PAS-B conformations generated by a WEMD simulation, as mentioned above. Consistent with our NOE-based ligand-receptor distances, the two top-scoring poses reveal two different orientations of KG-655 in the cavity. Furthermore, in each of these poses, the internal cavity is partially open to solvent (23% gain in solvent accessible surface area), and this opening appears to involve partial unfolding of the Fα helix in which the percentage of contacts that are present in the ARNT PAS-B crystal structure 4EQ1 has dropped to as little as 64%. Interestingly, we observed ARNT PAS-B precipitation when a high concentration (≥ 5 mM) of KG-655 was added. Precipitation was also seen when the ligand was added to the surface binding impaired Y456T mutant, consistent with internal binding requiring partial unfolding of the protein. Additionally, we also noticed substantial line broadening in ^15^N/^1^H HSQC spectra of ARNT PAS-B in the presence of KG-655 (**Fig. 4C**). Since the internal binding of the ligand is in the fast exchange regime, the line broadenings have likely resulted from partial unfolding, in agreement with the simulation data.

In closing, the importance of ARNT/coactivator interactions in mediating HIF and AHR activities are fundamental. Targeting ARNT PAS-B/CCC complexes with artificial inhibitors could provide a new avenue to modulate both pathways for therapeutic purposes. While the ligands used here may not be suitable for practical applications, our study has revealed two ‘hotspots’ on and within the ARNT PAS-B domain where ligands are likely to bind, laying the foundation for constructing novel therapeutic strategies. Furthermore, this work provides an excellent example of how NMR approaches can be used to readily characterize weak protein-ligand interactions.

## Data Availability

All data that support the findings of this study are available upon request.

## Supporting Materials

The Supplementary Information section contains the following information: Supplementary Tables S1 – S5; Figures S1 – S11.

## Author Contributions

XX, DCF, MLS, RS, LTC, and KHG designed the experiments; XX, JC, LPM, DCF, MLS, and RS performed the experiments; XX, JC, LPM, DCF, MLS, RS, LTC, and KHG analyzed the data and wrote the manuscript.

## Acknowledgments

The authors thank James Aramini (CUNY ASRC) for technical assistance, Guilherme Dal Poggetto for discussion on DOSY experiments, and members from the Gardner Laboratory for constructive comments. Several figures were created or modified with BioRender.com.

## References

1. Lu, H., Zhou, Q., He, J., Jiang, Z., Peng, C., Tong, R. et al. (2020) Recent advances in the development of protein-protein interactions modulators: mechanisms and clinical trials. Signal Transduct Target Ther 5, 213.

2. Modell, A. E., Blosser, S. L., and Arora, P. S. (2016) Systematic Targeting of Protein-Protein Interactions. Trends Pharmacol Sci 37, 702–713.

3. Henley, M. J., and Koehler, A. N. (2021) Advances in targeting ‘undruggable’ transcription factors with small molecules. Nat Rev Drug Discov 20, 669–688.

4. Wells, J. A., and McClendon, C. L. (2007) Reaching for high-hanging fruit in drug discovery at protein-protein interfaces. Nature 450, 1001–1009.

5. Chene, P. (2006) Drugs targeting protein-protein interactions. ChemMedChem 1, 400–411.

6. Jin, L., Wang, W., and Fang, G. (2014) Targeting protein-protein interaction by small molecules. Annu Rev Pharmacol Toxicol 54, 435–456.

7. Kolonko, M., and Greb-Markiewicz, B. (2019) bHLH-PAS Proteins: Their Structure and Intrinsic Disorder. Int J Mol Sci 20, 3653.

8. Mandl, M., and Depping, R. (2014) Hypoxia-inducible aryl hydrocarbon receptor nuclear translocator (ARNT) (HIF-1beta): is it a rare exception? Mol Med 20, 215–220.

9. Sorg, O. (2014) AhR signalling and dioxin toxicity. Toxicol Lett 230, 225–233.

10. Denison, M. S., and Nagy, S. R. (2003) Activation of the aryl hydrocarbon receptor by structurally diverse exogenous and endogenous chemicals. Annu Rev Pharmacol Toxicol 43, 309–334.

11. Safe, S., and Zhang, L. (2022) The Role of the Aryl Hydrocarbon Receptor (AhR) and Its Ligands in Breast Cancer. Cancers (Basel) 14, 5574.

12. Gruszczyk, J., Grandvuillemin, L., Lai-Kee-Him, J., Paloni, M., Savva, C. G., Germain, P. et al. (2022) Cryo-EM structure of the agonist-bound Hsp90-XAP2-AHR cytosolic complex. Nat Commun 13, 7010.

13. Tsuji, N., Fukuda, K., Nagata, Y., Okada, H., Haga, A., Hatakeyama, S. et al. (2014) The activation mechanism of the aryl hydrocarbon receptor (AhR) by molecular chaperone HSP90. FEBS Open Bio 4, 796–803.

14. Larigot, L., Juricek, L., Dairou, J., and Coumoul, X. (2018) AhR signaling pathways and regulatory functions. Biochim Open 7, 1–9.

15. Semenza, G. L. (1999) Regulation of mammalian O2 homeostasis by hypoxia-inducible factor 1. Annu Rev Cell Dev Biol 15, 551–578.

16. Semenza, G. L. (2012) Hypoxia-inducible factors in physiology and medicine. Cell 148, 399–408.

17. Majmundar, A. J., Wong, W. J., and Simon, M. C. (2010) Hypoxia-inducible factors and the response to hypoxic stress. Mol Cell 40, 294–309.

18. Burroughs, S. K., Kaluz, S., Wang, D., Wang, K., Van Meir, E. G., and Wang, B. (2013) Hypoxia inducible factor pathway inhibitors as anticancer therapeutics. Future Med Chem 5, 553–572.

19. Neavin, D. R., Liu, D., Ray, B., and Weinshilboum, R. M. (2018) The Role of the Aryl Hydrocarbon Receptor (AHR) in Immune and Inflammatory Diseases. Int J Mol Sci 19, 3851.

20. Bersten, D. C., Sullivan, A. E., Peet, D. J., and Whitelaw, M. L. (2013) bHLH-PAS proteins in cancer. Nat Rev Cancer 13, 827–841.

21. Kolluri, S. K., Jin, U. H., and Safe, S. (2017) Role of the aryl hydrocarbon receptor in carcinogenesis and potential as an anti-cancer drug target. Arch Toxicol 91, 2497–2513.

22. Hockel, M., and Vaupel, P. (2001) Tumor hypoxia: definitions and current clinical, biologic, and molecular aspects. J Natl Cancer Inst 93, 266–276.

23. Shah, Y. M., and Xie, L. (2014) Hypoxia-inducible factors link iron homeostasis and erythropoiesis. Gastroenterology 146, 630–642.

24. McGettrick, A. F., and O’Neill, L. A. J. (2020) The Role of HIF in Immunity and Inflammation. Cell Metab 32, 524–536.

25. Jun, J. C., Rathore, A., Younas, H., Gilkes, D., and Polotsky, V. Y. (2017) Hypoxia-Inducible Factors and Cancer. Curr Sleep Med Rep 3, 1–10.

26. Lv, Y., Zhao, S., Han, J., Zheng, L., Yang, Z., and Zhao, L. (2015) Hypoxia-inducible factor-1alpha induces multidrug resistance protein in colon cancer. Onco Targets Ther 8, 1941–1948.

27. Xia, Y., Choi, H. K., and Lee, K. (2012) Recent advances in hypoxia-inducible factor (HIF)-1 inhibitors. Eur J Med Chem 49, 24–40.

28. Koh, M. Y., Spivak-Kroizman, T., Venturini, S., Welsh, S., Williams, R. R., Kirkpatrick, D. L. et al. (2008) Molecular mechanisms for the activity of PX-478, an antitumor inhibitor of the hypoxia-inducible factor-1alpha. Mol Cancer Ther 7, 90–100.

29. Zimmer, M., Ebert, B. L., Neil, C., Brenner, K., Papaioannou, I., Melas, A. et al. (2008) Small-molecule inhibitors of HIF-2a translation link its 5’UTR iron-responsive element to oxygen sensing. Mol Cell 32, 838–848.

30. Welsh, S. J., Dale, A. G., Lombardo, C. M., Valentine, H., de la Fuente, M., Schatzlein, A., et al. (2013) Inhibition of the hypoxia-inducible factor pathway by a G-quadruplex binding small molecule. Sci Rep 3, 2799.

31. Albadari, N., Deng, S., and Li, W. (2019) The transcriptional factors HIF-1 and HIF-2 and their novel inhibitors in cancer therapy. Expert Opin Drug Discov 14, 667–682.

32. Fallah, J., and Rini, B. I. (2019) HIF Inhibitors: Status of Current Clinical Development. Curr Oncol Rep 21, 6.

33. Key, J., Scheuermann, T. H., Anderson, P. C., Daggett, V., and Gardner, K. H. (2009) Principles of ligand binding within a completely buried cavity in HIF2alpha PAS-B. J Am Chem Soc 131, 17647–17654.

34. Scheuermann, T. H., Li, Q., Ma, H. W., Key, J., Zhang, L., Chen, R. et al. (2013) Allosteric inhibition of hypoxia inducible factor-2 with small molecules. Nat Chem Biol 9, 271–276.

35. Scheuermann, T. H., Tomchick, D. R., Machius, M., Guo, Y., Bruick, R. K., and Gardner, K. H. (2009) Artificial ligand binding within the HIF2alpha PAS-B domain of the HIF2 transcription factor. Proc Natl Acad Sci U S A 106, 450–455.

36. Wu, D., Su, X., Lu, J., Li, S., Hood, B. L., Vasile, S. et al. (2019) Bidirectional modulation of HIF-2 activity through chemical ligands. Nat Chem Biol 15, 367–376.

37. Deeks, E. D. (2021) Belzutifan: First Approval. Drugs 81, 1921–1927.

38. Xu, R., Wang, K., Rizzi, J. P., Huang, H., Grina, J. A., Schlachter, S. T. et al. (2019) 3-[(1S,2S,3R)-2,3-Difluoro-1-hydroxy-7-methylsulfonylindan-4-yl]oxy-5-fluorobenzonitrile (PT2977), a Hypoxia-Inducible Factor 2alpha (HIF-2alpha) Inhibitor for the Treatment of Clear Cell Renal Cell Carcinoma. J Med Chem 62, 6876–6893.

39. Labrecque, M. P., Prefontaine, G. G., and Beischlag, T. V. (2013) The aryl hydrocarbon receptor nuclear translocator (ARNT) family of proteins: transcriptional modifiers with multi-functional protein interfaces. Curr Mol Med 13, 1047–1065.

40. Endler, A., Chen, L., and Shibasaki, F. (2014) Coactivator recruitment of AhR/ARNT1. Int J Mol Sci 15, 11100–11110.

41. Partch, C. L., and Gardner, K. H. (2011) Coactivators necessary for transcriptional output of the hypoxia inducible factor, HIF, are directly recruited by ARNT PAS-B. Proc Natl Acad Sci U S A 108, 7739–7744.

42. Ziello, J. E., Jovin, I. S., and Huang, Y. (2007) Hypoxia-Inducible Factor (HIF)-1 regulatory pathway and its potential for therapeutic intervention in malignancy and ischemia. Yale J Biol Med 80, 51–60.

43. Beischlag, T. V., Taylor, R. T., Rose, D. W., Yoon, D., Chen, Y., Lee, W. H. et al. (2004) Recruitment of thyroid hormone receptor/retinoblastoma-interacting protein 230 by the aryl hydrocarbon receptor nuclear translocator is required for the transcriptional response to both dioxin and hypoxia. J Biol Chem 279, 54620–54628.

44. Guo, Y., Scheuermann, T. H., Partch, C. L., Tomchick, D. R., and Gardner, K. H. (2015) Coiled-coil coactivators play a structural role mediating interactions in hypoxia-inducible factor heterodimerization. J Biol Chem 290, 7707–7721.

45. Kim, J. H., and Stallcup, M. R. (2004) Role of the coiled-coil coactivator (CoCoA) in aryl hydrocarbon receptor-mediated transcription. J Biol Chem 279, 49842–49848.

46. Partch, C. L., Card, P. B., Amezcua, C. A., and Gardner, K. H. (2009) Molecular basis of coiled coil coactivator recruitment by the aryl hydrocarbon receptor nuclear translocator (ARNT). J Biol Chem 284, 15184–15192.

47. Sadek, C. M., Jalaguier, S., Feeney, E. P., Aitola, M., Damdimopoulos, A. E., Pelto-Huikko, M. et al. (2000) Isolation and characterization of AINT: a novel ARNT interacting protein expressed during murine embryonic development. Mech Dev 97, 13–26.

48. Guo, Y., Partch, C. L., Key, J., Card, P. B., Pashkov, V., Patel, A. et al. (2013) Regulating the ARNT/TACC3 axis: multiple approaches to manipulating protein/protein interactions with small molecules. ACS Chem Biol 8, 626–635.

49. Gagné, D., Azad, R., Edupuganti, U. R., Williams, J., Aramini, J. M., Akasaka, K., et al. (2020) Use of High Pressure NMR Spectroscopy to Rapidly Identify Proteins with Internal Ligand-Binding Voids. bioRxiv 2020.2008.2025.267195.

50. Ma, B., Elkayam, T., Wolfson, H., and Nussinov, R. (2003) Protein-protein interactions: structurally conserved residues distinguish between binding sites and exposed protein surfaces. Proc Natl Acad Sci U S A 100, 5772–5777.

51. Sheffield, P., Garrard, S., and Derewenda, Z. (1999) Overcoming expression and purification problems of RhoGDI using a family of “parallel” expression vectors. Protein Expr Purif 15, 34–39.

52. Senn, H., Werner, B., Messerle, B. A., Weber, C., Traber, R., and Wüthrich, K. (1989) Stereospecific assignment of the methyl 1H NMR lines of valine and leucine in polypeptides by nonrandom 13C labelling. FEBS Letters 249, 113–118.

53. Card, P. B., Erbel, P. J. A., and Gardner, K. H. (2005) Structural basis of ARNT PAS-B dimerization: Use of a common beta-sheet interface for hetero– and homodimerization. J Mol Biol 353, 664–677.

54. Breeze, A. L. (2000) Isotope-filtered NMR methods for the study of biomolecular structure and interactions. Progress in Nuclear Magnetic Resonance Spectroscopy 36, 323–372.

55. Iwahara, J., Wojciak, J. M., and Clubb, R. T. (2001) Improved NMR spectra of a protein– DNA complex through rational mutagenesis and the application of a sensitivity optimized isotope-filtered NOESY experiment. Journal of Biomolecular NMR 19, 231–241.

56. Ogura, K., Terasawa, H., and Inagaki, F. (1996) An improved double-tuned and isotope-filtered pulse scheme based on a pulsed field gradient and a wide-band inversion shaped pulse. J Biomol NMR 8, 492–498.

57. Zwahlen, C., Legault, P., Vincent, S. J. F., Greenblatt, J., Konrat, R., and Kay, L. E. (1997) Methods for Measurement of Intermolecular NOEs by Multinuclear NMR Spectroscopy: Application to a Bacteriophage λ N-Peptide/boxB RNA Complex. Journal of the American Chemical Society 119, 6711–6721.

58. Neri, D., Szyperski, T., Otting, G., Senn, H., and Wuthrich, K. (1989) Stereospecific nuclear magnetic resonance assignments of the methyl groups of valine and leucine in the DNA-binding domain of the 434 repressor by biosynthetically directed fractional ^13^C labeling. Biochemistry 28, 7510–7516.

59. Szyperski, T., Neri, D., Leiting, B., Otting, G., and Wuthrich, K. (1992) Support of 1H NMR assignments in proteins by biosynthetically directed fractional 13C-labeling. J Biomol NMR 2, 323–334.

60. Johnson, B. A. (2018) From Raw Data to Protein Backbone Chemical Shifts Using NMRFx Processing and NMRViewJ Analysis. Methods Mol Biol 1688, 257–310.

61. Norris, M., Fetler, B., Marchant, J., and Johnson, B. A. (2016) NMRFx Processor: a cross-platform NMR data processing program. J Biomol NMR 65, 205–216.

62. Morris, K. F., and Johnson, C. S., Jr. (1993) Resolution of discrete and continuous molecular size distributions by means of diffusion-ordered 2D NMR spectroscopy. Journal of the American Chemical Society 115, 4291–4299.

63. Williamson, M. P. (2013) Using chemical shift perturbation to characterise ligand binding. Prog Nucl Magn Reson Spectrosc 73, 1–16.

64. Russo, J. D., Zhang, S., Leung, J. M. G., Bogetti, A. T., Thompson, J. P., DeGrave, A. J. et al. (2022) WESTPA 2.0: High-performance upgrades for weighted ensemble simulations and analysis of longer-timescale applications. Journal of Chemical Theory and Computation 18, 638–649.

65. Case, D. A., Belfon, K., Ben-Shalom, I. Y., Brozell, S. R., Cerutti, D. S., Cheatham, I., T. E., et al. (2020) AMBER University of California, San Francisco

66. Williams, C. J., Headd, J. J., Moriarty, N. W., Prisant, M. G., Videau, L. L., Deis, L. N. et al. (2018) MolProbity: More and better reference data for improved all-atom structure validation. Protein Sci 27, 293–315.

67. Tian, C., Kasavajhala, K., Belfon, K. A. A., Raguette, L., Huang, H., Migues, A. N. et al. (2020) ff19SB: Amino-Acid-Specific Protein Backbone Parameters Trained against Quantum Mechanics Energy Surfaces in Solution. J Chem Theory Comput 16, 528–552.

68. Izadi, S., Anandakrishnan, R., and Onufriev, A. V. (2014) Building Water Models: A Different Approach. J Phys Chem Lett 5, 3863–3871.

69. Ryckaert, J. P., Ciccotti, G., and Berendsen, H. J. C. (1977) Numerical integration of the cartesian equations of motion of a system with constraints: molecular dynamics of n-alkanes. J Comput Phys 23, 327–341.

70. Essmann, U., Perera, L., Berkowitz, M. L., Darden, T., Lee, H., and Pedersen, L. G. (1995) A smooth particle mesh ewald method. J Chem Phys 103, 8577–8593.

71. Torrillo, P. A., Bogetti, A. T., and Chong, L. T. (2021) A Minimal, Adaptive Binning Scheme for Weighted Ensemble Simulations. J Phys Chem A 125, 1642–1649.

72. Gonzalez-Aleman, R., Hernandez-Castillo, D., Caballero, J., and Montero-Cabrera, L. A. (2020) Quality Threshold Clustering of Molecular Dynamics: A Word of Caution. J Chem Inf Model 60, 467–472.

73. Dominguez, C., Boelens, R., and Bonvin, A. M. (2003) HADDOCK: a protein-protein docking approach based on biochemical or biophysical information. J Am Chem Soc 125, 1731–1737.

74. Jorgensen, W. L., and Tirado-Rives, J. (1988) The OPLS [optimized potentials for liquid simulations] potential functions for proteins, energy minimizations for crystals of cyclic peptides and crambin. J Am Chem Soc 110, 1657–1666.

75. He, X., Man, V. H., Yang, W., Lee, T. S., and Wang, J. (2020) A fast and high-quality charge model for the next generation general AMBER force field. J Chem Phys 153, 114502.

76. Joung, I. S., and Cheatham, T. E., 3rd. (2008) Determination of alkali and halide monovalent ion parameters for use in explicitly solvated biomolecular simulations. J Phys Chem B 112, 9020–9041.

77. Bayly, C. I., Cieplak, P., Cornell, W. D., and Kollman, P. A. (1993) A well-behaved electrostatic potential based method using charge restraints for deriving atomic charges: The RESP Model. Journal of Physical Chemistry 97, 10269–10280.

78. Krouglova, T., Vercammen, J., and Engelborghs, Y. (2004) Correct diffusion coefficients of proteins in fluorescence correlation spectroscopy. Application to tubulin oligomers induced by Mg2+ and Paclitaxel. Biophys J 87, 2635–2646.

79. Huber, G. A., and Kim, S. (1996) Weighted-ensemble Brownian dynamics simulations for protein association reactions. Biophysical Journal 70, 97–110.

80. Zuckerman, D. M., and Chong, L. T. (2017) Weighted ensemble simulation: Review of methodology, applications, and software. Ann Rev Biophys 46, 43–57.

81. Sztain, T., Ahn, S. H., Bogetti, A. T., Casalino, L., Goldsmith, J. A., Seitz, E. et al. (2021) A glycan gate controls opening of the SARS-CoV-2 spike protein. Nature Chemistry 13, 963–968.

82. Saglam, A. S., and Chong, L. T. (2019) Protein-protein binding pathways and calculations of rate constants using fully-continuous, explicit-solvent simulations. Chemical Sciences 10, 2360.

83. Zwier, M. C., Pratt, A. J., Adelman, J. L., Kaus, J. W., Zuckerman, D. M., and Chong, L. T. (2016) Efficient atomistic simulation of pathways and calculation of rate constants for a protein-peptide binding process: Application to the MDM2 protein and an intrinsically disordered p53 peptide. J Phys Chem Lett 7, 3440–3445.

84. Brossard, E. E., and Corcelli, S. A. (2023) Molecular Mechanism of Ligand Binding to the Minor Groove of DNA. J Phys Chem Lett 14, 4583–4590.

85. Evans, M. R., Card, P. B., and Gardner, K. H. (2009) ARNT PAS-B has a fragile native state structure with an alternative beta-sheet register nearby in sequence space. Proc Natl Acad Sci U S A 106, 2617–2622.

86. Evans, M. R., and Gardner, K. H. (2009) Slow transition between two beta-strand registers is dictated by protein unfolding. J Am Chem Soc 131, 11306–11307.

87. Xu, X., Dikiy, I., Evans, M. R., Marcelino, L. P., and Gardner, K. H. (2021) Fragile protein folds: Sequence and environmental factors affecting the equilibrium of two interconverting, stably folded protein conformations. Magn Reson (Gott) 2, 63–76.

88. Wilkins, S. E., Abboud, M. I., Hancock, R. L., and Schofield, C. J. (2016) Targeting Protein–Protein Interactions in the HIF System. ChemMedChem 11, 773–786.

89. Dai, S., Qu, L., Li, J., Zhang, Y., Jiang, L., Wei, H. et al. (2022) Structural insight into the ligand binding mechanism of aryl hydrocarbon receptor. Nat Commun 13, 6234.

90. Henry, J. T., and Crosson, S. (2011) Ligand-binding PAS domains in a genomic, cellular, and structural context. Annu Rev Microbiol 65, 261–286.

91. Mangraviti, A., Raghavan, T., Volpin, F., Skuli, N., Gullotti, D., Zhou, J. et al. (2017) HIF-1alpha-Targeting Acriflavine Provides Long Term Survival and Radiological Tumor Response in Brain Cancer Therapy. Sci Rep 7, 14978.

92. Wu, D., Potluri, N., Lu, J., Kim, Y., and Rastinejad, F. (2015) Structural integration in hypoxia-inducible factors. Nature 524, 303–308.

93. Ren, X., Diao, X., Zhuang, J., and Wu, D. (2022) Structural basis for the allosteric inhibition of hypoxia-inducible factor (HIF)-2 by belzutifan. Mol Pharmacol

