## Supplementary Information for "Identification of Small Molecule Ligand Binding Sites On and In the ARNT PAS-B Domain"

Contents:
Tables S1-S5
Figures S1-S11

**Table S1. NOE Assignments for ARNT PAS-B/KG-655**

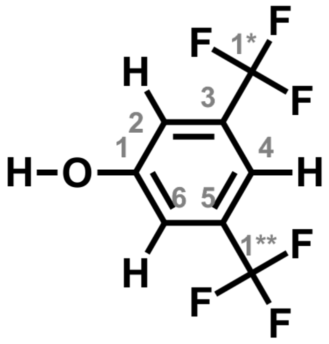


3,5-Bis(trifluoromethyl)phenol

| Protein Assignment | Ligand Assignment | Distance Range (Å) |
|---|---|---|
| I396HD11 | H4 | 1.8-5.0 |
| I396HD11 | H2/H6 | 1.8-5.0 |
| I396HG21 | H2/H6 | 1.8-2.8 |
| # I458HG21 | H4 | 1.8-2.8 |
| I457HG21 | H4 | 1.8-3.5 |
| V425HG11 | H2/H6 | 1.8-3.5 |
| * V381HG11 | H4 | 1.8-3.5 |
| V415HG11 | H4 | 1.8-3.5 |
| L418HD21 | H4 | 1.8-5.0 |
| L418HD21 | H2/H6 | 1.8-3.5 |
| L408HD11 | H2/H6 | 1.8-2.8 |
| V415HG21 | H4 | 1.8-5.0 |
| L423HD11 | H2/H6 | 1.8-3.5 |
| L418HG | H2/H6 | 1.8-3.5 |

Surface-facing residues marked with #. Outliers (do not belong to either binding site) are marked with *.

**Table S2. NOE Assignments for ARNT PAS-B/KG-279**

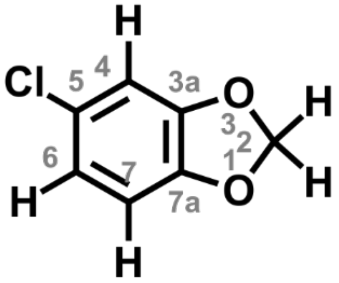

5-chloro-1,3-benzodioxole

| Protein Assignment | Ligand Assignment | Distance Range (Å) |
|---|---|---|
| I396HD11 | H4 | 1.8-5.0 |
| I396HD11 | H6 | 1.8-3.5 |
| I396HD11 | H7 | 1.8-3.5 |
| I457HD11 | H4 | 1.8-5.0 |
| I457HD11 | H6 | 1.8-5.0 |
| I457HD11 | H7 | 1.8-5.0 |
| I457HD11 | H2a/H2b | 1.8-2.8 |
| I396HG21 | H4 | 1.8-3.5 |
| I396HG21 | H6 | 1.8-3.5 |
| I396HG21 | H7 | 1.8-2.8 |
| # I458HG21 | H4 | 1.8-5.0 |
| # I458HG21 | H6 | 1.8-5.0 |
| # I458HG21 | H7 | 1.8-5.0 |
| I457HG21 | H4 | 1.8-5.0 |
| I457HG21 | H6 | 1.8-5.0 |
| I457HG21 | H7 | 1.8-5.0 |
| I457HG21 | H2a/H2b | 1.8-5.0 |
| V425HG21 | H4 | 1.8-2.8 |
| V425HG21 | H6 | 1.8-3.5 |
| V425HG21 | H7 | 1.8-5.0 |
| V425HG21 | H2a/H2b | 1.8-5.0 |
| L418HD21 | H4 | 1.8-3.5 |
| L418HD21 | H6 | 1.8-2.8 |
| L418HD21 | H7 | 1.8-3.5 |
| L408HD11 | H4 | 1.8-3.5 |
| L408HD11 | H6 | 1.8-3.5 |
| L408HD11 | H7 | 1.8-5.0 |
| V415HG21 | H4 | 1.8-3.5 |
| L408HD21 | H4 | 1.8-5.0 |
| L408HD21 | H6 | 1.8-5.0 |
| L408HG | H4 | 1.8-3.5 |

Surface-facing residues marked with #.

**Table S3. GAFF2 atomic charges derived for KG-655 drug ligand.** Atom names in the structure of the ligand correspond to those used for ligand parameterization. Note that these atom names differ from labels by IPUAC locants in Tables S1 and S2.

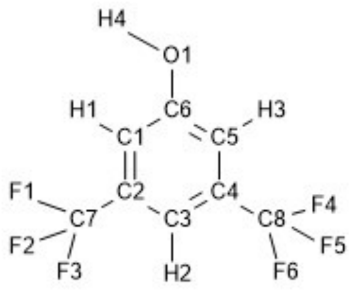


| Atom Name | Atom Type | Partial Atomic Charges |
|---|---|---|
| C1/C5 | CA | -0.344626 |
| H1/H3 | HA | 0.208731 |
| C2/C4 | CA | -0.020023 |
| C7/C8 | C3 | 0.660231 |
| F1/F2/F3 | F | -0.214117 |
| C3 | CA | -0.193060 |
| H2 | HA | 0.167975 |
| F4/F5/F6 | F | -0.214117 |
| C6 | CA | 0.458901 |
| O1 | OH | -0.549457 |
| H4 | HO | 0.391717 |

**Table S4. GAFF2 atomic charges derived for KG-279 drug ligand.** Atom names in the structure of the ligand correspond to those used for ligand parameterization. Note that these atom names differ from labels by IPUAC locants in Tables S1 and S2.

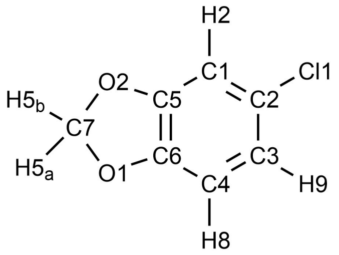


| Atom Name | Atom Type | Partial Atomic Charges |
| --- | --- | --- |
| C1 | CA | -0.238490 |
| H2 | HA | 0.203597 |
| C5 | CA | 0.287911 |
| C2 | CA | -0.067733 |
| Cl1 | CL | -0.101533 |
| C3 | CA | -0.042623 |
| H9 | HA | 0.150656 |
| C4 | CA | -0.431844 |
| H8 | HA | 0.239701 |
| C6 | CA | 0.384687 |
| O1 | OS | -0.447371 |
| C7 | C3 | 0.388408 |
| H5a/H5b | H2 | 0.055106 |
| O2 | OS | -0.435578 |

**Table S5. Diffusion coefficients of ARNT PAS-B in the presence of KG-548, 655, and 279**

| | | | |
|---|---|---|---|
| | **100μM ARNT PAS-B** | | |
| | **Diffusion Coefficients** | **$D:D_{monomer}$** | **%monomer** |
| No Ligand | $1.14\times10^{-6}$ $cm^2/sec$ | 1.00 | 100% |
| 5mM KG-548 | $9.19\times10^{-7}$ $cm^2/sec$ | 0.81 | 24% |
| 5mM KG-655 | $9.55\times10^{-7}$ $cm^2/sec$ | 0.84 | 41% |
| 2mM KG-279 | $9.92\times10^{-7}$ $cm^2/sec$ (adjusted) | 0.87 | 57% |
| | **280μM ARNT PAS-B** | | |
| | **Diffusion Coefficients** | **$D:D_{monomer}$** | **%monomer** |
| No Ligand | $1.06\times10^{-6}$ $cm^2/sec$ | 0.93 | 80% |
| 5mM KG-548 | $9.01\times10^{-7}$ $cm^2/sec$ | 0.79 | 14% |
| 5mM KG-655 | $9.28\times10^{-7}$ $cm^2/sec$ | 0.82 | 29% |
| 2mM KG-279 | $1.00\times10^{-6}$ $cm^2/sec$ (adjusted) | 0.88 | 60% |
| | **600μM ARNT PAS-B** | | |
| | **Diffusion Coefficients** | **$D:D_{monomer}$** | **%monomer** |
| No Ligand | $9.89\times10^{-7}$ $cm^2/sec$ | 0.87 | 56% |
| 5mM KG-548 | $9.10\times10^{-7}$ $cm^2/sec$ | 0.80 | 19% |
| 5mM KG-655 | $9.00\times10^{-7}$ $cm^2/sec$ | 0.79 | 14% |
| 2mM KG-279 | $9.43\times10^{-7}$ $cm^2/sec$ (adjusted) | 0.83 | 38% |

| | | |
|---|---|---|
| | **Example Water Diffusion Coefficients** | |
| | **100μM ARNT PAS-B** | **600μM ARNT PAS-B** |
| No Ligand | $2.41\times10^{-5}$ $cm^2/sec$ | $2.34\times10^{-5}$ $cm^2/sec$ |
| 5mM KG-548 | $2.36\times10^{-5}$ $cm^2/sec$ | $2.37\times10^{-5}$ $cm^2/sec$ |
| 5mM KG-655 | $2.41\times10^{-5}$ $cm^2/sec$ | $2.34\times10^{-5}$ $cm^2/sec$ |
| 2mM KG-279 | $1.79\times10^{-5}$ $cm^2/sec$ | $1.75\times10^{-5}$ $cm^2/sec$ |

Diffusion coefficients of ARNT PAS-B were calculated using the integrations at varying gradient strengths of the aliphatic region between 0.5-1 ppm. Diffusion coefficients were used to determine the degree of dimerization in each sample, such that double the molecular weight would correspond to a diffusion coefficient that is 79.8% of the monomer diffusion coefficient (1). Apo ARNT PAS-B was assumed to be 100% monomeric at 100μM. Dimerization of ARNT PAS-B was found to occur at both high concentrations as well as in the presence of small molecule ligand, with KG-548 having the most drastic effect. Diffusion coefficients in the presence of KG-279 were adjusted based on water diffusion coefficients measurements.

### SI Figures

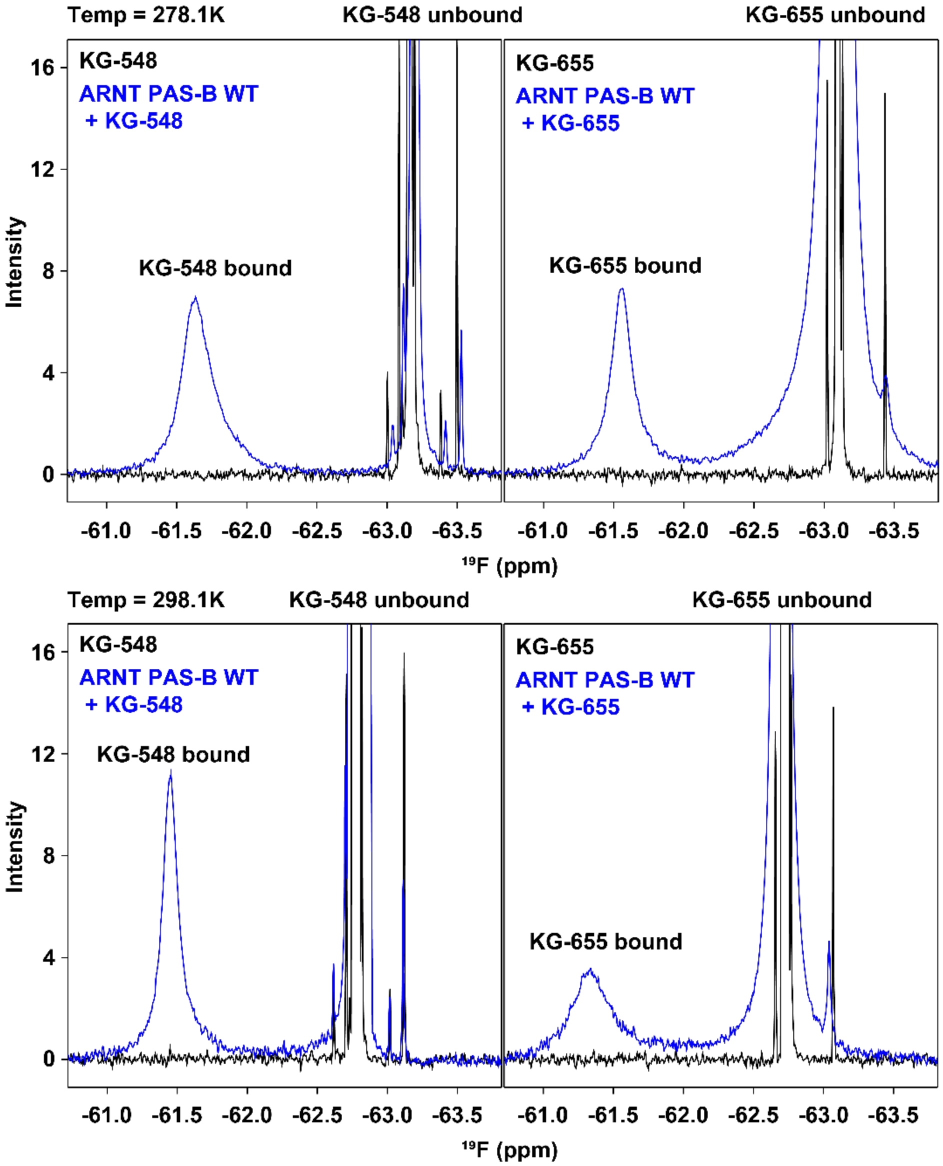


**Figure S1. Evaluation of KG-548 and KG-655 ligand interactions with ARNT PAS-B using 1D $^{19}$F NMR at 278K.1 and 298.1K.**

The surface-bound spectra of both ligands showed an additional peak between -61 and -61.5 ppm. The free ligand peak of KG-655 was also broadened in the presence of ARNT PAS-B, indicating a possible second binding mode.

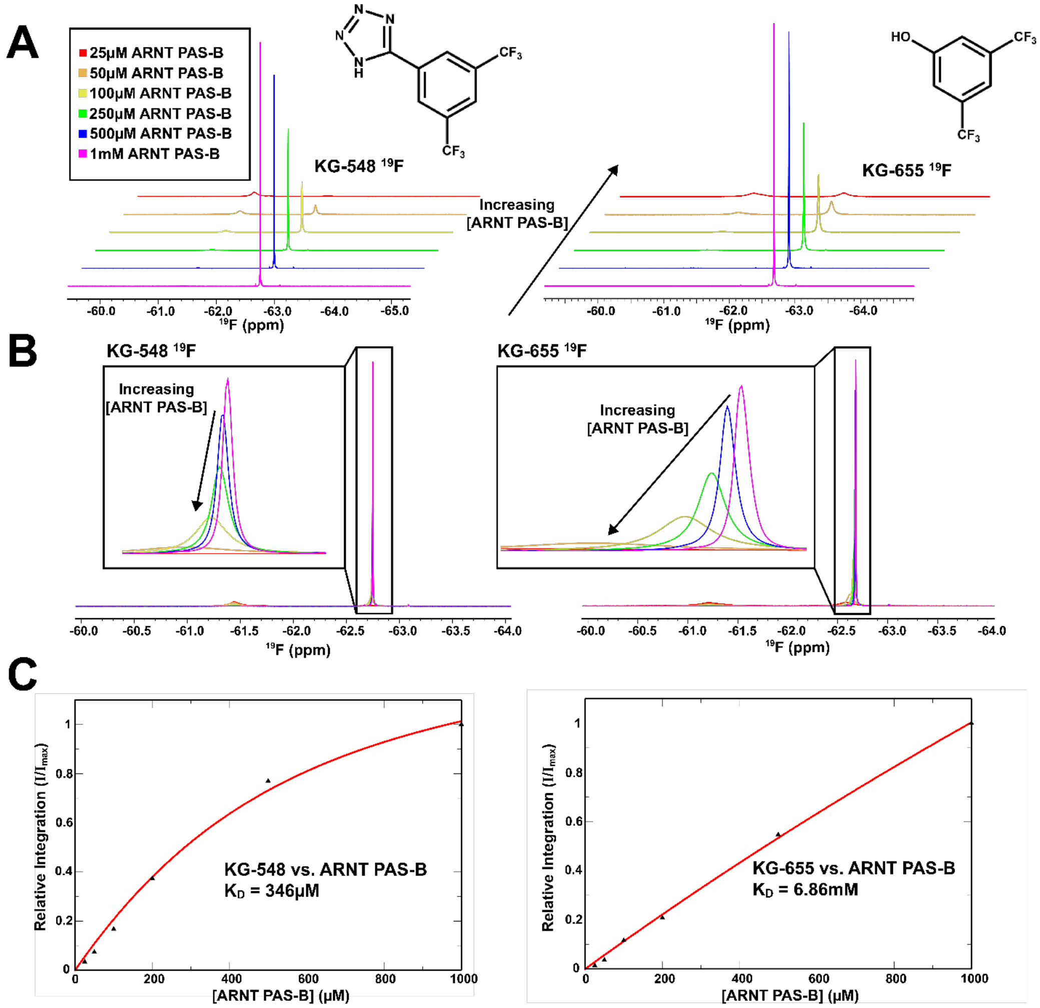


**Figure S2. Evaluating the Binding Affinity of KG-548 and KG-655 via $^{19}$F NMR.**
A) Stacked $^{19}$F spectra in the presence of either KG-548 (left) or KG-655 (right) with varying concentrations of ARNT PAS-B. As ARNT PAS-B concentration increases, there is a decrease in the intensity of a peak at 62.7 ppm, complemented by the appearance of a second peak at 61.3 ppm. B) The same $^{19}$F spectra as in panel A but superimposed to show a downfield shift in the peak at 62.7 ppm as the concentration of ARNT PAS-B increases. This shift is noticeably more drastic in KG-655 than in KG-548, possibly due to the existence of a second binding mode that is in the fast-exchange regime. C) The relative integration of the peaks at 61.3 ppm plotted against the concentration of ARNT PAS-B (left: KG-548, right: KG-655). We note that the KG-548 shape of the KG-548 titration data show deviations from the ideal one-site binding curve used to fit the data, which we attribute to the ARNT PAS-B homodimerization event discussed elsewhere in the text. We have not attempted to account for this monomer:dimer equilibrium in our fitting here, so the $K_D$ = 346μM value should be treated as an estimate. Similarly, the lack of saturable binding and the two binding modes of KG-655 complicate obtaining a precise $K_D$ value; we show here a fit to a single-binding mode model but underscore the value itself should be treated as an estimate.

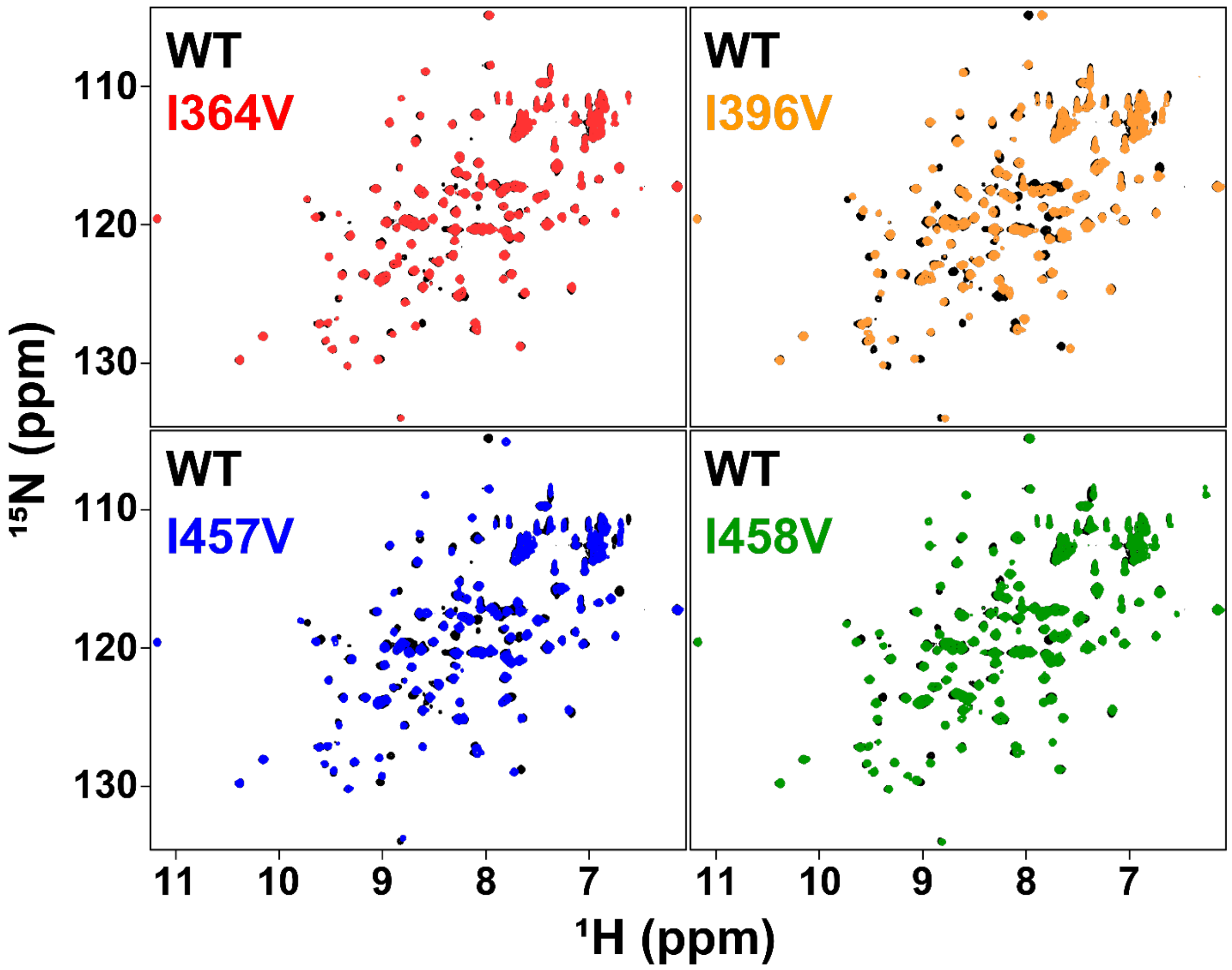


**Figure S3. Comparisons of the four Ile to Val mutants to the WT ARNT PAS-B protein.** $^{15}$N/$^{1}$H-HSQC spectra of the four Ile to Val ARNT PAS-B mutants (250 µM) indicated these mutants were folded and adopted similar structures to the WT protein, as evident by good chemical shift dispersion and overlap with the spectrum of the WT ARNT PAS-B.

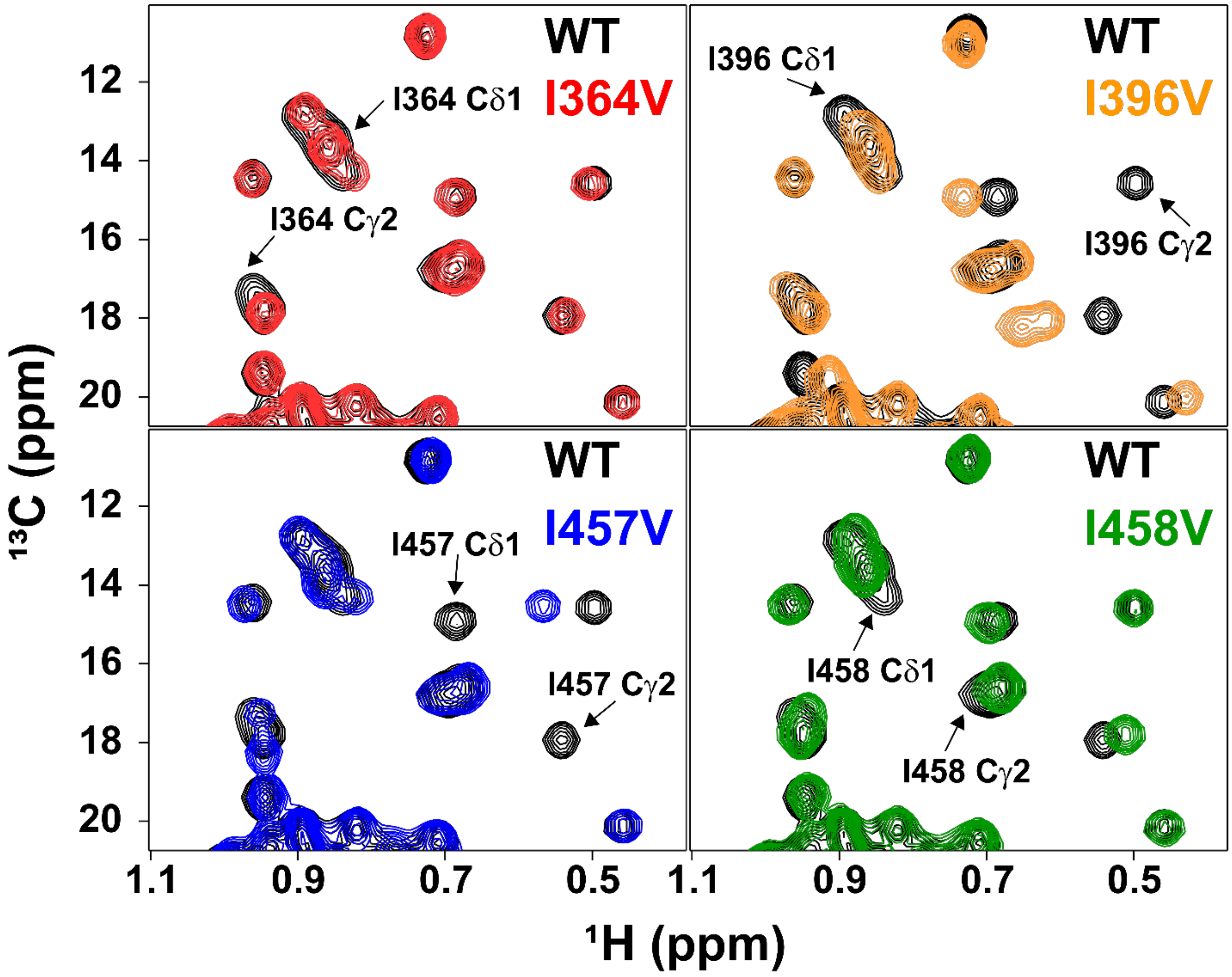


**Figure S4. $^{13}C/^{1}H$-HSQC spectra of the ARNT PAS-B mutants compared with the wildtype (WT) protein.**

$^{13}C/^{1}H$-HSQC spectra of the four Ile to Val ARNT PAS-B mutants (250 µM) were used to confirm the Ile methyl assignments. The methyl cross-peak locations of the mutated residues are marked with arrows. Peaks other than the mutated Ile residues showed good overlap with the spectrum of the WT protein.

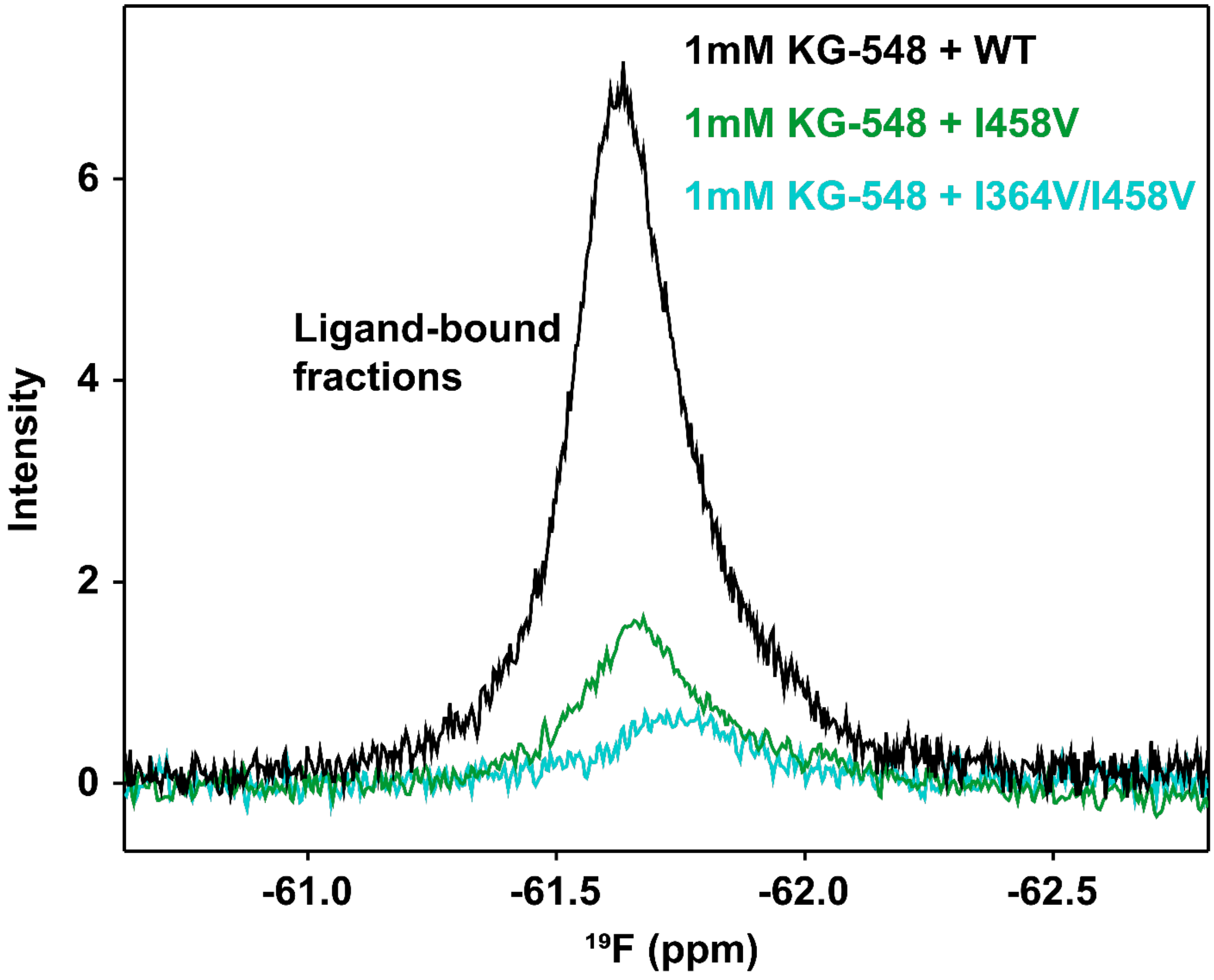


**Figure S5. I364V/I458V double mutation further disrupts surface binding.**
$^{19}$F-spectra of 1 mM KG-548 mixed with 250 µM ARNT PAS-B WT, I458V, or I364V/I458V double mutant, zoomed in on the bound-fraction of the ligand. Mutation I458V substantially decreased binding affinity, with the I364V/I458V double mutation resulting in the lowest amount of bound ligand.

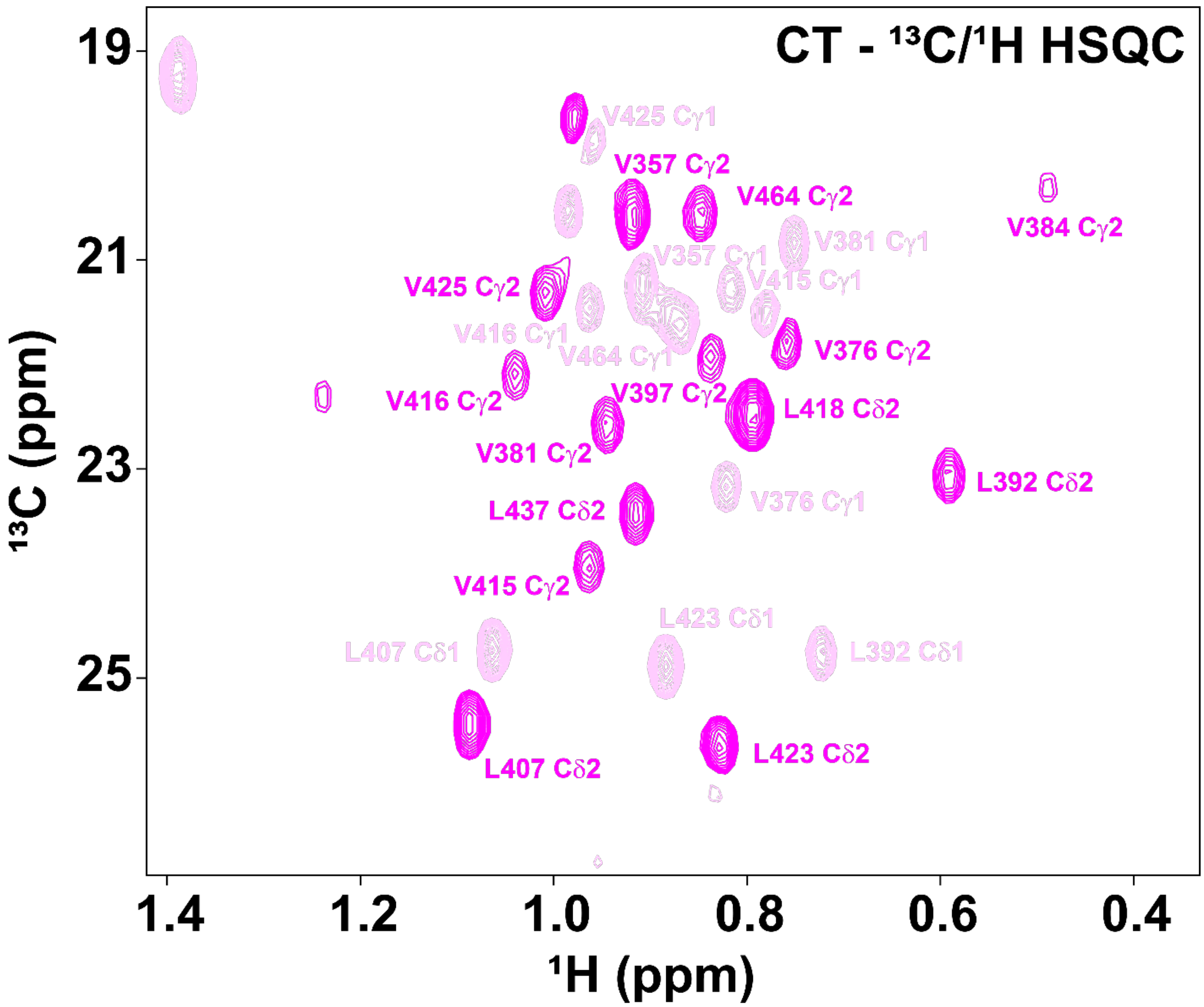


**Figure S6. Stereospecific assignments of Leu and Val methyl resonances.**
Pro-R methyl groups (γ1 and δ1 for Val and Leu, respectively) yield negative signals in this constant time (CT) -$^{13}C/^{1}H$ HSQC experiment. Pro-S methyl groups (γ2 and δ2 for Val and Leu, respectively) yield positive signals. Pro-S methyl groups are the migrating methyl groups. When they are $^{13}C$ labeled, the adjoining carbons are highly unlikely also to be labeled. Signs of the signals are determined by checking the signs of the Met methyl peaks, which are positives here.

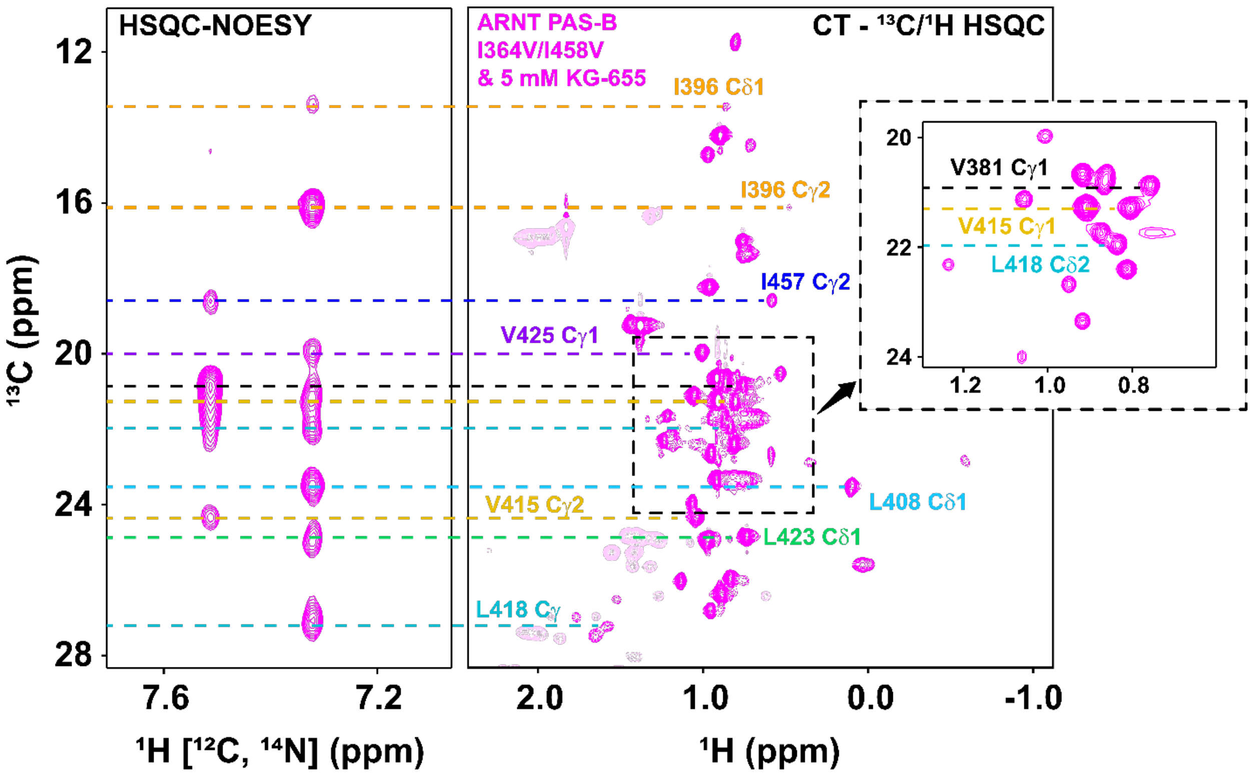


**Figure S7. The surface and internal binding modes of KG-655 are independent of each other.**

A double-filtered HSQC-NOESY experiment was performed using KG-655 and ARNT PAS-B I364V/I458V double mutant. Heteronuclear NOE correlations between KG-655 and methyl groups of all of the internal-facing protein residues identified previously using the WT ARNT PAS-B remain unaffected.

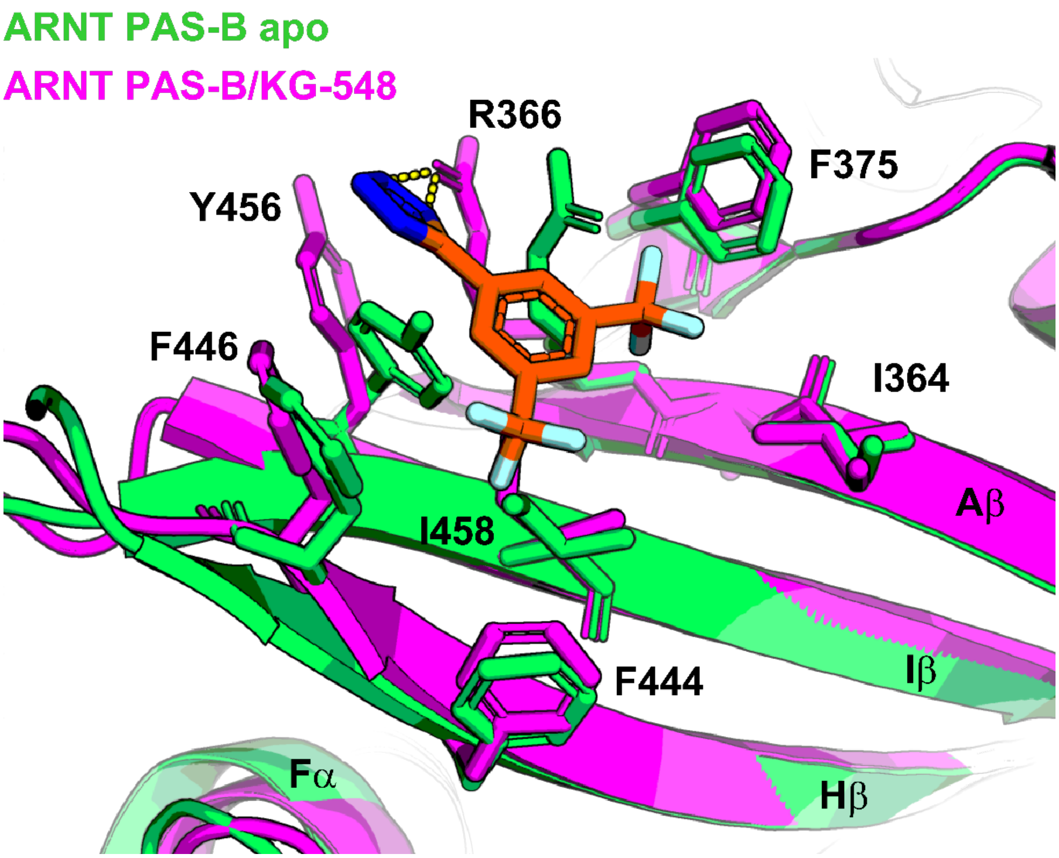


**Figure S8. Comparison between ARNT PAS-B apo and KG-548 bound states.**
Superimposition of the crystal structure of the ARNT PAS-B apo (PDB: 4EQ1, lime green) and KG-548 bound state (PDB: 8G4A, magenta), with residues within 5Å of the ligand shown in sticks. Residue Y456 moves away from KG-548 in the ligand-bound state to prevent clashes. R366 also reorients and forms polar contacts with the tetrazole group of KG-548 (yellow dashed lines).

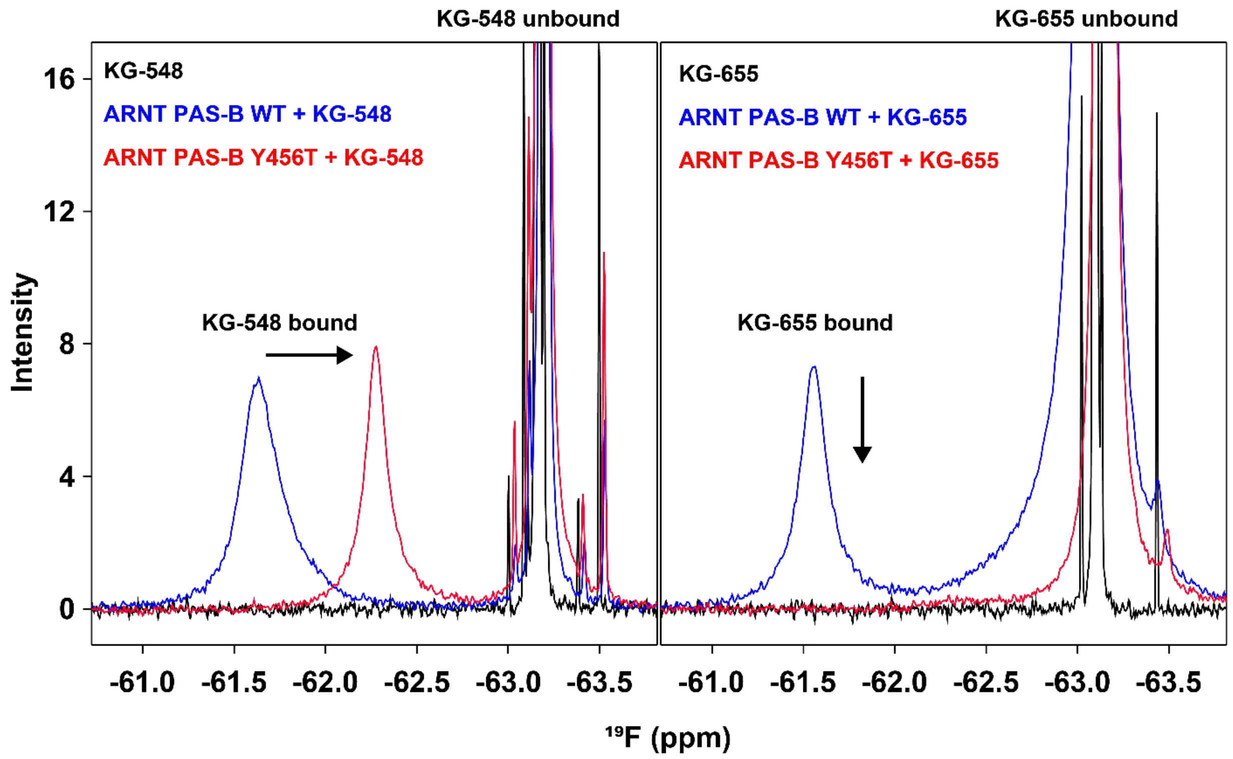


**Figure S9. Mutation Y456T disrupts the surface binding of KG-655.**
$^{19}$F spectra of ligands KG-548 or KG-655 (1 mM) mixed with ARNT PAS-B WT and Y456T (250 µM), collected at 278.1K. Mutation Y456T affected the surface binding of the two ligands differently, shifting the ligand-bound peak of KG-548 upfield by 0.65 ppm while completely abolishing the surface binding of KG-655. The internal binding mode of KG-655 also appeared to be affected (less broadening of the peak at -63.1 ppm) by the mutation.

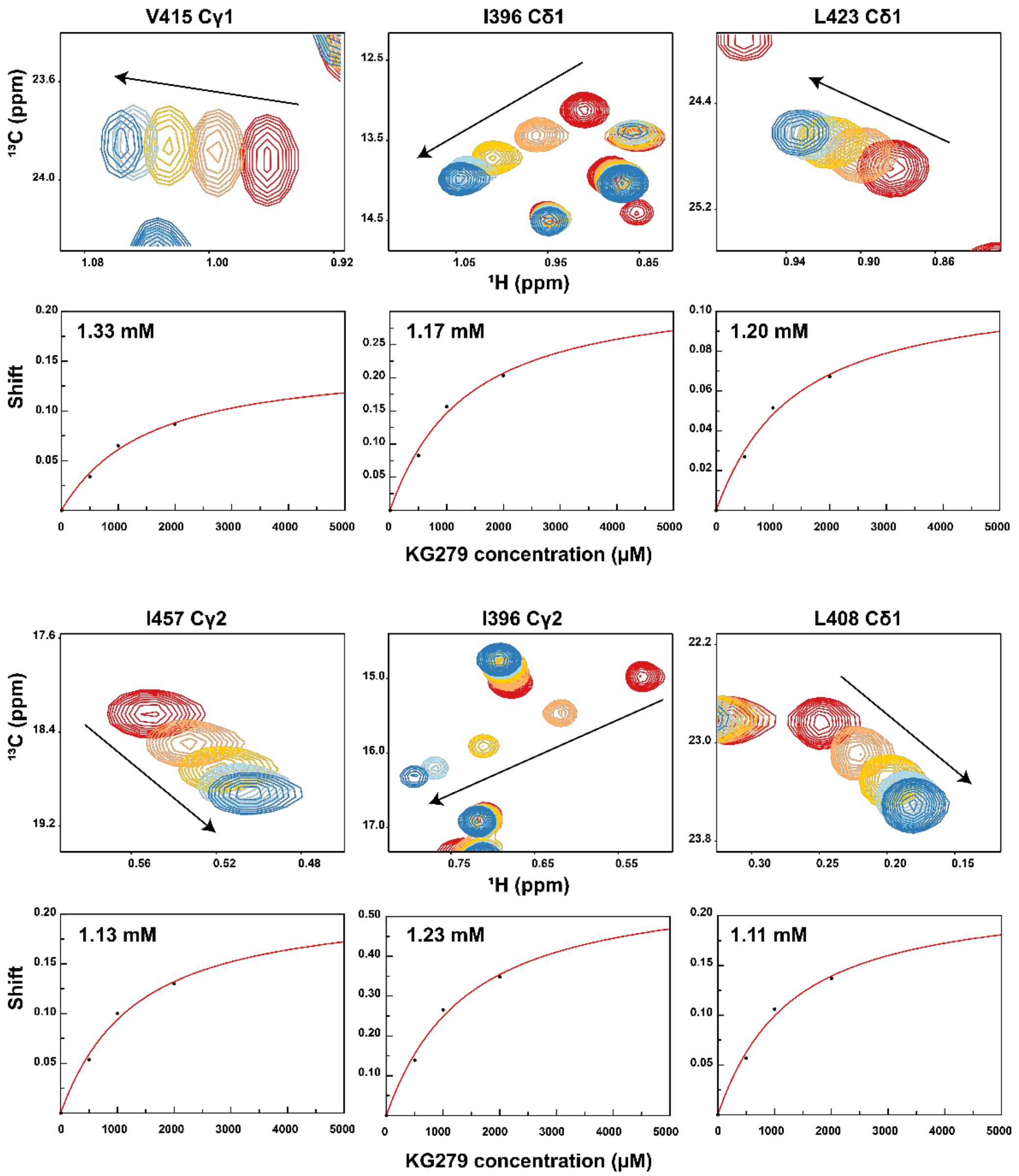


**Figure S10. Evaluating the binding affinity of KG-279 through ligand titration into ARNT PAS-B.** Six methyl peaks showing the largest ligand-dependent chemical shift changes were utilized to estimate the binding affinity of KG-279. Contour plots of these peaks are zoomed-in views of the constant time (CT)-$^{13}$C/$^{1}$H HSQC spectra from Figure 6B in the main text. The titration experiment included five concentrations (0, 500, 1000, 2000, and 5000 μM). The highest concentration (5000 μM) was excluded from the fitting analysis due to protein precipitation observed post data acquisition.

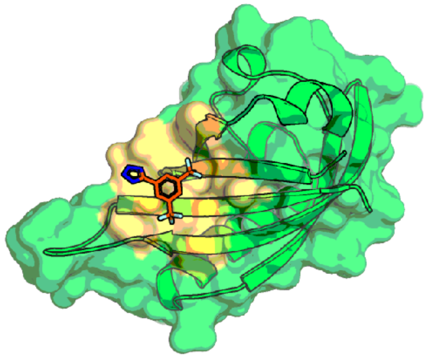

ARNT PAS-B / KG-548

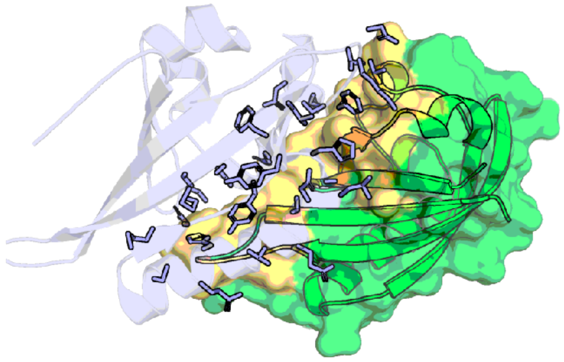

ARNT PAS-B / HIF-2α

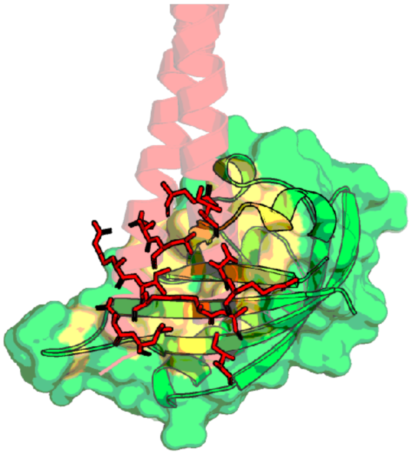

ARNT PAS-B / hTACC3

**Figure S11. The ARNT PAS-B/KG-548 interface is an important surface binding 'hotspot'.** The same β-sheet surface has also been shown to interact with HIF-2α PAS-B and TACC3 coactivator.

1. Krouglova, T., Vercammen, J., and Engelborghs, Y. (2004) Correct diffusion coefficients of proteins in fluorescence correlation spectroscopy. Application to tubulin oligomers induced by Mg2+ and Paclitaxel. *Biophys J* **87**, 2635-2646
